# Thalamus orchestrates local acetylcholine-dependent dopamine release in the learning striatum

**DOI:** 10.64898/2026.05.08.723861

**Authors:** Andrew J. Miller-Hansen, ManHua Zhu, Ryan F. Kovaleski, Baran Demir, Talia N. Lerner

## Abstract

Dopamine is essential for striatal function and learning. Striatal dopamine release can be triggered by dopamine cell firing, but also by coordinated cholinergic interneuron activity, which stimulates dopamine release via presynaptic nicotinic acetylcholine receptors on dopamine axons. While acetylcholine-dependent dopamine release is well-documented *ex vivo* and under artificial optogenetic stimulation *in vivo*, its role during natural behavior has remained unclear. One possible endogenous driver of acetylcholine-dependent dopamine release is thalamic input, which provides strong excitatory drive to cholinergic interneurons. To examine whether thalamic input provokes acetylcholine-dependent dopamine release during behavior, we performed simultaneous fiber photometry recordings of striatal dopamine (GRAB-rDA3m) and thalamic axon activity (gCaMP8m) in the dorsomedial (DMS) and dorsolateral striatum (DLS) of mice learning the accelerating rotarod, a striatal-dependent task that demands precise and effortful motor control. Recordings were obtained on- and off-task and across days of training to capture the full arc of learning. Dopamine transients in DMS, but not DLS, were frequently coupled to peaks in thalamic axon activity via an acetylcholine-dependent mechanism. The occurrence of these thalamic-evoked DMS dopamine transients depended on learning, task engagement, and the recent history of dopamine activity, but did not contribute to motor error signals. Together, these findings establish thalamic input as a physiological driver of acetylcholine-dependent dopamine release in DMS. Moreover, they reveal that striatal sensitivity to this local release mechanism is dynamically gated by dopaminergic history, providing a compelling framework for understanding how local and soma-triggered dopamine signals are coordinated to support learning.

## MAIN

Dopamine (**DA**) is a critical neurotransmitter that governs learning, motivation, and movement^1,2^, but the mechanisms underlying behaviorally relevant DA release remain debated^3^. While foundational studies of DA have highlighted the patterns of action potentials fired by cell bodies in the midbrain^4,5^, which are consistent with these neurons’ computational role in reward prediction errors^6,7^, other studies indicate that DA release can be evoked in the striatum independently of DA cell body firing via local mechanisms at striatal terminals^8–10^. Acetylcholine (**ACh**) released by striatal cholinergic interneurons (**ChIs**) depolarizes DA axons – sometimes even evoking action potentials – and initiates neurotransmitter release, independently of DA cell bodies *ex vivo*^9,11^. However, it is controversial whether this phenomenon occurs under physiological conditions during behavior^3,12^, especially since ACh and DA are frequently anti-correlated *in vivo*^13,14^. Furthermore, it is unclear how local cell-body-independent and soma-triggered dopamine release mechanisms might be coordinated to shape striatal function. Notably, recent studies in the dorsal striatum of rodents have failed to find evidence for physiological ACh-dependent DA release either during spontaneous behavior^13^ or a 2-armed bandit task^15^. However, given other reports of ACh-dependent DA release supporting effortful behavior in the nucleus accumbens in rodents^16^ and song maturation in juvenile songbirds^17^, we hypothesized that ACh-dependent DA release might occur in the dorsal striatum over the course of learning an effortful motor task, and that observations of ACh-dependent DA release *in vivo* might depend on specific conditions where the microcircuit is receptive to such a mechanism.

## Thalamic input precedes DA release in DMS but not DLS during accelerating rotarod learning

To explore the possibility that ACh-dependent DA release occurs during an effortful motor task, we used the accelerating rotarod, a striatum-dependent motor learning task^18,19^ that is free from explicit rewards (which evoke canonical reward responses in midbrain DA cell bodies). To probe physiological mechanisms that might induce task-dependent dorsal striatal DA release via ChI activation, we considered known natural drivers of ChIs. Thalamic inputs to the striatum provide particularly strong excitatory drive to ChIs^20–23^, and optogenetic stimulation of thalamostriatal axons can induce ACh-dependent DA release in brain slices^8–10^. To examine the relationship between thalamostriatal activity and DA release during motor learning, we performed dual-color *in vivo* fiber photometry recordings. We simultaneously measured the activity of thalamic axons and DA release in the dorsomedial striatum (**DMS**) and dorsolateral striatum (**DLS**) across stages of motor learning (Fig. 1a, S1a).

**Fig. 1:**
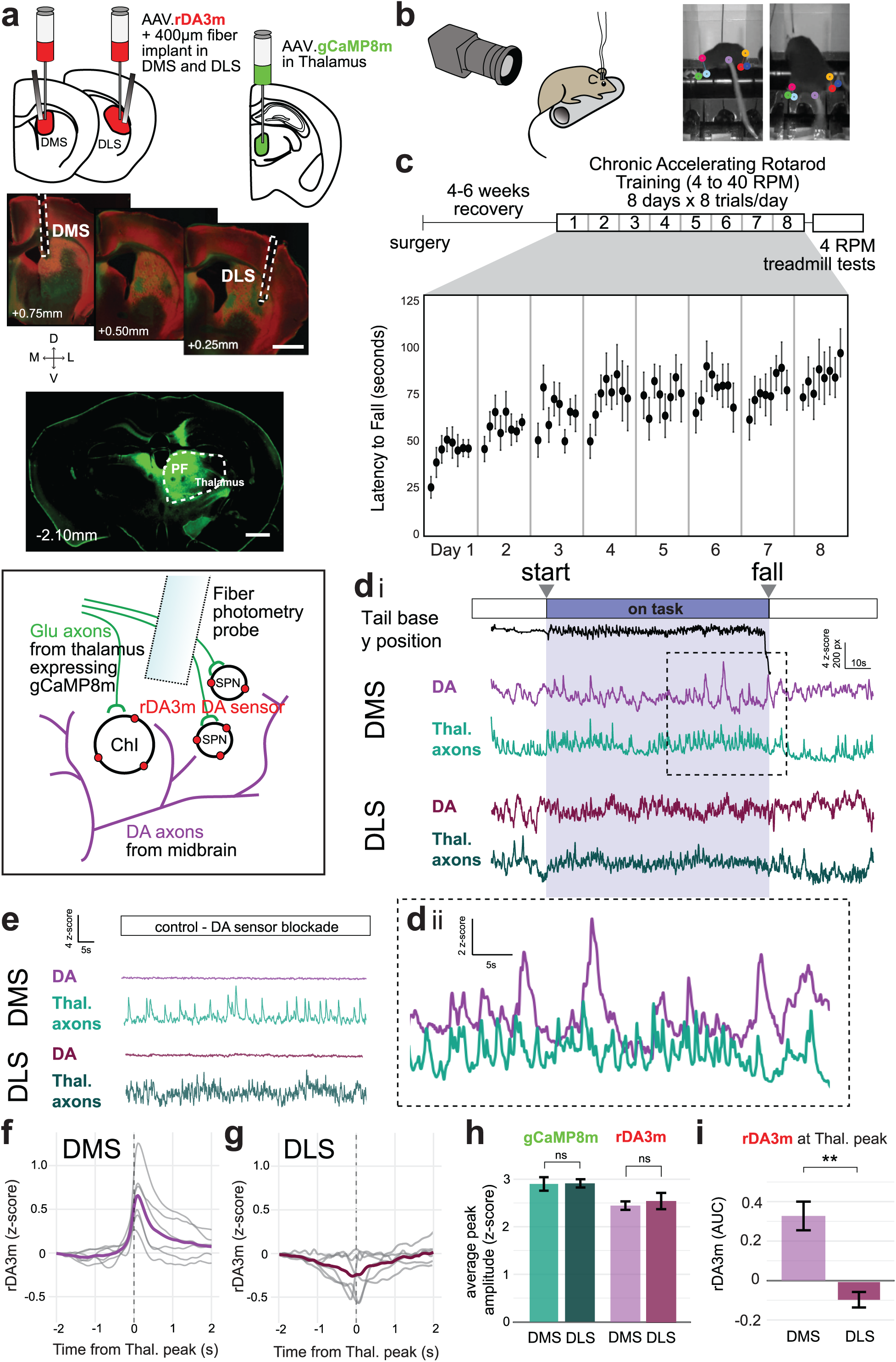
Thalamic-linked DA release events occur in DMS but not DLS during accelerating rotarod learning. a, Schematic of viral expression strategy for simultaneous measurement of DA release and thalamic axon activity in both DMS and DLS via *in vivo* fiber photometry. Scale bars 1mm. Representative image of thalamic gCaMP8m expression, showing thalamus (dotted line) and parafasicular (PF) nucleus, where injections were centered. b, Video tracking of hind limbs and tail during accelerated rotarod learning. c, Chronic accelerated rotarod training schedule and average latency to fall for animals in Fig. 1 (n=9) over 8 days of training. d, Representative trace of a rotarod trial showing tracked tail base position with simultaneously collected thalamic axon (gCaMP8m) activity and DA release (rDA3m) in DMS and DLS. Inset dii shows a close-up of the gCaMP8m and rDA3m fluctuations co-activating in DMS. e, The rDA3m signal was dependent on the binding of DA, and was flattened by blockade of the D1R-based sensor with the D1R antagonist SCH-23390 (SCH, 10mg/kg), without affecting the gCaMP8m signal. f, rDA3m signal during thalamic axon gCaMP8m transients in DMS. All mice showed notable DA increases during these moments (gray lines, averages from individual animals, light purple line, average across n=6 mice). g, The same as f, but in DLS. DA increases were absent during thalamic input to DLS, rather some mice showed decreases in DA release on average during these moments (same 6 mice as in f). h, The amplitudes of gCaMP8m transients in DMS vs. DLS (averages from n=6 mice, p=0.810 ns, MWU test) and amplitudes of rDA3m transients in DMS vs. DLS (averages from n=6 mice, p=0.471 ns, MWU test). i, rDA3m signal area under curve (AUC) during thalamic gCaMP8m transients in DMS vs. DLS (averages from n=6 mice, p=0.005** MWU test). Barplots throughout show mean with +/- standard error bars.

We virally expressed the calcium sensor gCaMP8m^26^ in the thalamus (AAV9-Syn-gCaMP8m), targeting an area centered on the parafasicular nucleus (Fig. 1a, S1b), whose axons densely innervate all subregions of dorsal striatum^20,28^. We virally expressed the red-shifted DA sensor GRAB-rDA3m^25^ in DMS and DLS (AAV2/9-hSyn-rDA3m). We then implanted 400 μm-diameter fiber-optic probes in DMS and DLS for dual-color recordings in each region. After recovery from surgery, mice began chronic accelerating rotarod training for 8 days x 8 trials/day with *in vivo* fiber photometry and paw and tail tracking (Fig. 1b-c). Rotarod trials began with ∼30s standing stationary on the rod before it started rotating, accelerating from 4 to 40RPM (“on task”, Fig. 1di) over 300s or until the subject fell. Photometry data were collected for an additional ∼30 seconds after each fall. Importantly, calcium activity of thalamic axons could be measured reliably independently of DA release, without cross-channel interference or hemodynamic artifacts, as indicated by pharmacological controls that blocked the dopamine-binding site of the rDA3m sensor and did not alter the calcium signal (Fig. 1e).

DMS, but not DLS, showed robust co-activation of thalamic axon activity and DA release (Fig. 1dii, f, g, i). DMS DA transients lagged DMS thalamic activity by ∼100-150ms (Fig. 1f). Co-activation was absent in DLS signals recorded in the same mice, ruling out behavioral variability as an explanation (Fig. 1g). Co-activation in DMS was absent when sham viruses (EYFP and mCherry) were expressed in place of gCaMP8m and rDA3m, during DA sensor block (Fig. 1e), and after time-shifting the gCaMP8m signal (Fig. S1c), confirming that the co-activation is related to neural activity and temporally specific. The prominent coactivation in DMS compared to DLS was not due to stronger thalamic signals in this subregion, since the amplitudes of thalamic axon peaks were not significantly different between DMS and DLS (Fig. 1h). Likewise, although the DA signal specifically during concurrent thalamic axon activity was much larger in DMS than DLS, there were no overall differences in DA peak amplitudes between DMS and DLS (Fig. 1h-i).

## Thalamic input to DMS evokes dopamine via a nAChR-dependent mechanism

Peaks in thalamic axon activity reliably *preceded* DA peaks in DMS, consistent with a proposed mechanism of ChI activation and nicotinic acetylcholine receptor (**nAChR**)-dependent depolarization of DA terminals. Therefore, we tested whether pharmacological blockade of nAChRs with mecamylamine (Mec.; 3mg/kg i.p., or saline control) would specifically prevent thalamus-associated DA events. Because nicotinic blockers can have anti-kinetic effects^29^, which might indirectly alter the striatal dynamics under investigation, we tested mice on a slow (4 RPM) rotarod “treadmill test” to allow for a recording session where mice remained moving at a consistent speed pre- and post-drug delivery (Fig. 2a). At 4 RPM, most mice had no difficulty remaining on the rotarod without falling, even after Mec. treatment. One mouse fell repeatedly during treadmill testing, and its data were excluded from the subsequent analyses.

**Fig. 2:**
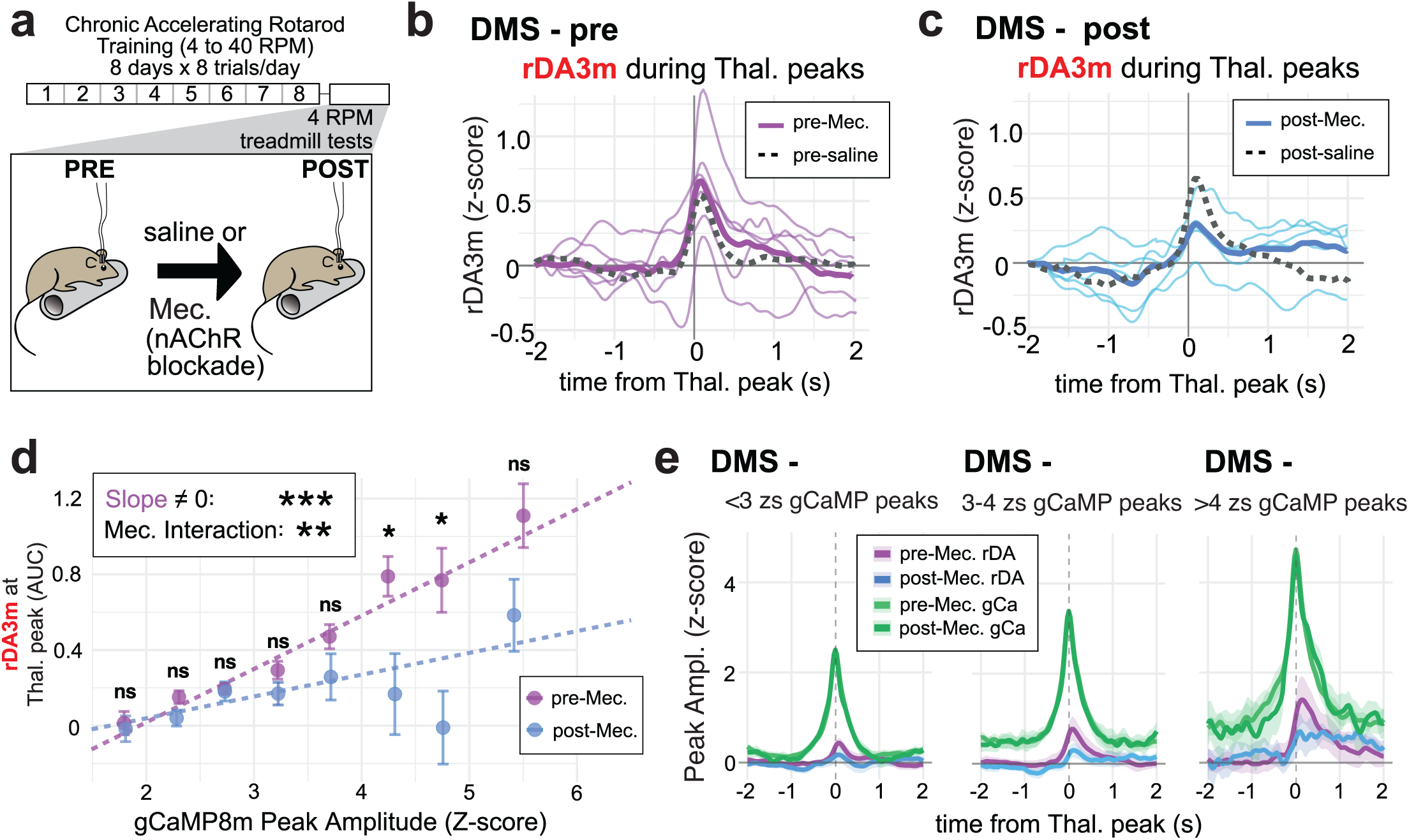
Thalamic input to DMS evokes DA via a nAChR-dependent mechanism. a, Experimental schedule showing trials 1-8 and off-task recordings in between trials. b, rDA3m responses during thalamic gCaMP8m peaks before Mec. (thin lines represent individual averages, thick line is the average from all mice). Dotted gray line is the average from pre-saline recordings (n=6 mice). c, The same as b, but after MEC injection (n=5 mice). Dotted gray line is post-saline average (n=5 mice). d, Binned gCaMP8m peaks in DMS and their concurrent rDA3m responses. There is a significant effect of gCaMP8m peak amplitude on rDA3m response (linear regression interaction model, see methods for details, slope=0.269 for pre-Mec., p<0.001***), and this relationship was significantly altered by Mec. (slope=0.093 for post-Mec., Mec. interaction, Fisher Z test p=0.009**). Bins were 0.5 Z-scores wide, and unpaired Wilcoxon Rank-Sum tests were done within each bin, with the following results for all 8 bins: 0.788ns, 0.097ns, 0.648ns, 0.116ns, 0.093ns, 0.015*, 0.021*, 0.069ns. e, Average thalamic gCamp8m peaks and concurrent rDA3m responses during small (<3 Zs), moderate (3-4), and large (>4) gCaMP8m events. Average gCaMP8m peaks pre/post Mec. overlap.

Neither Mec. nor saline administration affected overall thalamic axon peak amplitudes (Fig. S2a) or overall DA peak amplitudes in DMS or DLS (Fig. S2b), but Mec. treatment appeared to reduce thalamus-associated DMS DA responses by nearly half (Fig. 2b-c, Fig. S2c), although this effect did not reach significance at the animal level (DMS p=0.106; DLS p=0.590). However, because widespread synchrony of ChI firing is required to elicit nAChR-dependent DA release^8,27^, we reasoned that larger thalamic axon peaks representing the strongest or most synchronized thalamic input might more strongly elicit DA peaks sensitive to nAChR blockade. Indeed, when we binned thalamic axon peak amplitudes, we found that Mec. significantly reduced coincident DA responses during large thalamic events (>4 Z-score, Fig. 2d-e). This analysis also established that thalamic axon peak amplitude itself correlated positively with the magnitude of the concurrent DA response, and the slope of this relationship was significantly altered by Mec. (Fig. 2d). In DLS, as during the rotarod training, there was no meaningful co-activation of thalamic activity and DA release, and Mec. had no effect (Fig. S2c,e). Because Mec. specifically reduced DMS DA release coincident with strong thalamic input - without significantly altering overall DA release - we concluded that thalamic activity evokes DMS DA release via an ACh-dependent mechanism *in vivo* during the observed gCaMP8m/rDA3m co-activations. Residual DA release observed during nAChR blockade is consistent with two recent *in vivo* pharmacological studies of locally-evoked DA via ChI activation, which reported similar partial effects of nicotinic block^17^, and identified muscarinic and other receptors that contribute to DA release evoked by ChI stimulation^27^.

## Thalamus-evoked DA release in DMS changes over the course of training while thalamic activation of ChIs remains stable

Interleaved between rotarod trial recordings were “off-task” recordings, when the mice moved spontaneously or sat in the bottom well of the rotarod device for 2 minutes before the next rotarod trial started (Fig. 3a). These off-task recordings allowed us to determine the features of gCaMP8m/rDA3m signals that were specific to task performance, as well as how signals changed over the course of training. In DLS, both thalamic peaks and DA peaks were slightly larger during task performance and grew over training (Fig. S3a-b), but there was no change over training in DA at the time of thalamic input: the correlation remained slightly negative throughout training (Fig. 3b, S3c). In DMS, thalamic peaks and DA peaks were also slightly larger during task performance but only DA peaks grew over training, peaking on the last day, Day 8 (Fig. S3d-e). Thalamus-evoked DMS DA peaks increased significantly over days of rotarod performance, peaking on Day 6 (Fig. 3c-e, S3f), despite stable amplitudes of thalamic input (Fig. S3d).

**Fig. 3:**
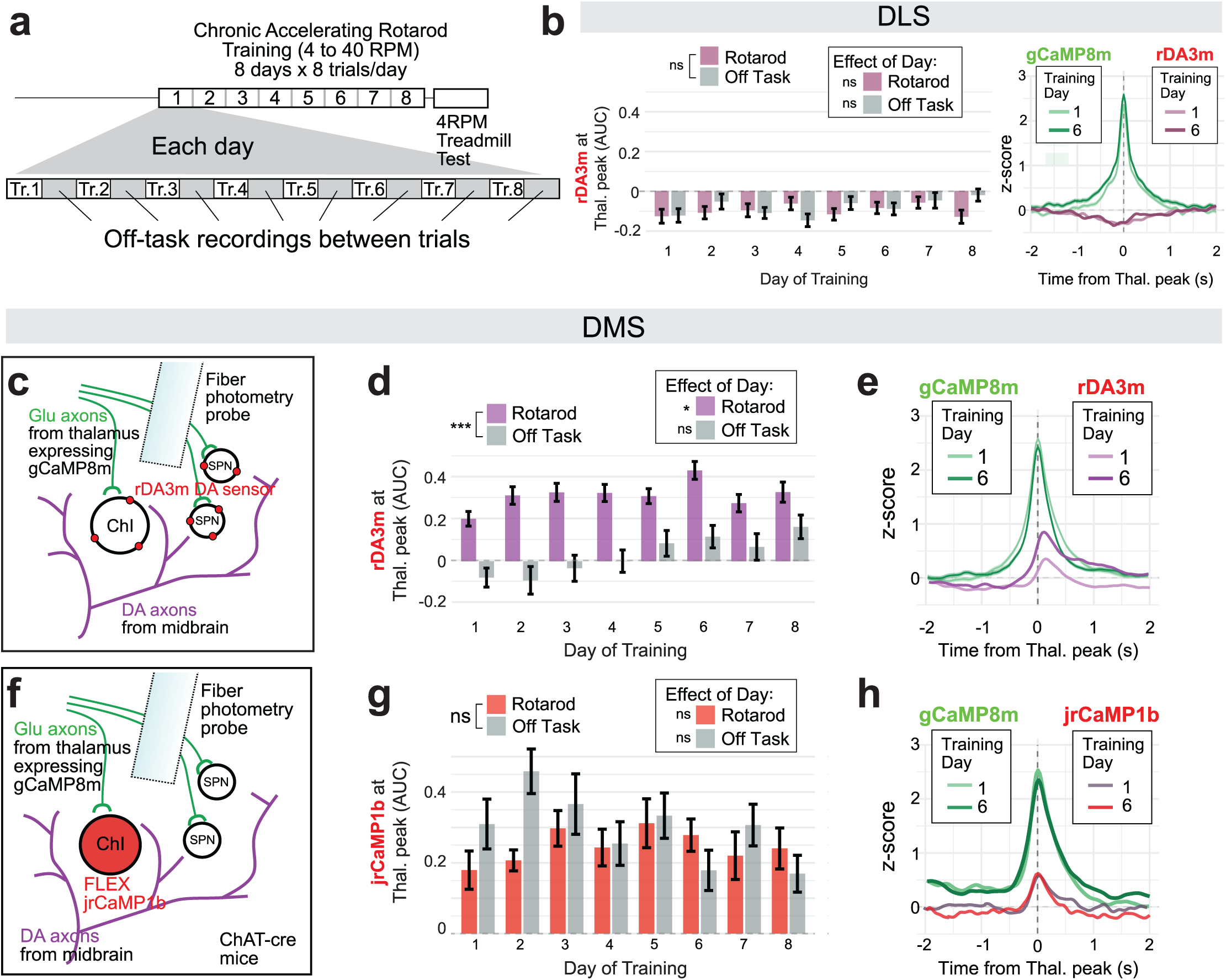
Thalamic-linked DA release events are task-dependent and increase during learning but thalamic ChI activation does not. a, Experimental schedule. b, AUC quantifications of rDA3m response during thalamic gCaMP8m peaks in DLS, showing no effect of training day during rotarod trials (F(7, 34.36) = 0.80, p = 0.595ns) or off-task recordings (F(7, 35.23) = 1.12, p = 0.371ns). Rotarod trials and off-task recordings showed similar values overall (F(1, 709.32) = 1.30, p = 0.254ns). Traces (right) show averaged gCaMPm peak on day 1 vs. 6 and averaged concurrent rDA3m responses. c, Diagram of sensor expression strategy for dual-color rDA3m/gCaMP8m experiments. d, AUC quantifications as in b, but in DMS. Significant effect of day on rDA3m signal in DMS for rotarod trials (LMM: F(7, 34.49) = 2.61, p = 0.029*) but not for off-task recordings (F(7, 35.02) = 2.22, p = 0.056ns), as well as a significant difference between on and off-task conditions (F(1, 708.85) = 198.76, p < 0.001***). e, Traces show averaged gCaMP8m peak on day 1 vs. day 6 and averaged concurrent rDA3m responses in DMS. f, Diagram of sensor expression strategy for dual-color jrCaMP1b/gCaMP8m experiments in DMS only. g, jrCaMP1b signal during thalamic peaks over days of training (AUC quantified as in rDA3m experiments, n=3 mice), no significant effect of day for rotarod trials (LMM: F(7,14.00)=0.37, p=0.907ns) or off-task recordings (F(7,14.03)=2.10, p=0.113 ns), and no significant difference between behavioral conditions (F(1,357.07)=3.77, p=0.053 ns). h, Traces show averaged gCaMP8m peak on day 1 vs. 6 and concurrent jrCaMP1b responses in DMS.

Changes in thalamus-evoked DA release could result from learning-related plasticity or from faster locomotion, as subjects remain on the rotarod longer and reach higher rotation speeds on later days of training. However, the amplitude of thalamus-evoked DA events was not significantly correlated with rotarod speed (Fig. S4a). We also compared thalamus-evoked DA only during the first 45s of each trial. Mice generally stay on the rotarod for at least 45s, even on Day 1 of training, and these first 45s are performed at the same speed regardless of training day. In this analysis, we still observed increases in thalamus-evoked DMS DA over days of training, also peaking on Day 6 (Fig. S4b), suggesting the learning-related increase is independent of increased locomotion.

If synchronized thalamic input to DMS is evoking local DA release via synchronous activation of ChIs, we reasoned that we should also be able to observe thalamic activation of ChIs at short latency *in vivo*. Therefore, we expressed the red-shifted calcium sensor jrCaMP1b in ChIs (AAV1-CAG-FLEX-NES-jRCaMP1b injection into ChAT-IRES-cre mice), and gCaMP8m in thalamus as in previous experiments (Fig. 3f). We trained these mice on the same chronic accelerating rotarod task for 8 days while recording dual-color fiber photometry in DMS, and observed ChI activity peaks that lagged behind thalamic peaks, as predicted, but with a shorter latency than DA peaks (∼40ms compared to 100-150ms observed for DA). Interestingly, however, there was no clear change in the strength of thalamic-evoked ChI activation over days, either during rotarod performance or in off-task recordings, and no obvious task dependence (Fig. 3g-h, S5a-b), contrasting with the observed task-dependence and changes over learning in the thalamus-evoked DA signal (Fig 3d-e vs. g-h), and suggesting that learning-related plasticity and task-dependent gating of thalamus-evoked DA occurs downstream of the thalamus-ChI synapse.

To confirm our *in vivo* findings and to more closely evaluate whether the properties of thalamostriatal synapses onto ChIs could contribute to the region-specific (DMS>DLS) relationship between thalamic input and DA release, we performed optogenetics-assisted slice physiology experiments (Fig. S5c-d). We expressed ChR2-EYFP in thalamus (AAV9-hSyn-hChR2(H134R)-EYFP), allowing us to optogenetically activate thalamic inputs and verify functional thalamostriatal synapses in striatal brain slices. To target ChIs for recording, we crossed ChAT-IRES-cre mice to an Ai14 reporter line to express tdTomato in ChIs. We prepared acute slices for synaptic physiology and recorded excitatory postsynaptic currents (EPSCs) in tdTomato-identified ChIs in DMS and DLS while optogenetically activating thalamic ChR2+ axons with a blue LED (1ms, ∼5mW/mm²). Evoked responses in DMS and DLS ChIs showed characteristics previously associated with thalamus-ChI synapses, including fast rise times, large NMDAR currents, and constitutive AMPAR rectification^22,30,31^. There were no significant differences between thalamic synapses onto ChIs in DMS and DLS on any measured variable (EPSC amplitude, AMPA:NMDA, Paired-Pulse Ratio (PPR), rise time, Fig. S5e-h), except for AMPAR rectification index, which was slightly increased in DMS (Fig. S5i). These data demonstrate that the machinery for ChI engagement by excitatory thalamic inputs is equivalent in DMS and DLS and cannot explain region-dependent differences in thalamus-linked DA release events observed *in vivo*.

## Thalamus Does Not Drive DA Error Signals in Striatum

Since strong thalamostriatal input evokes DMS DA during rotarod performance, we examined this relationship more closely to evaluate its potential computational function. DA is classically associated with reward prediction error, a computational function well suited to support learning^6^. Analogous action prediction errors in dorsal striatum have been suggested to support motor learning^32^. While the rotarod task contains no explicit reward, one highly salient error signal occurs when mice fall from the rotarod at the end of each trial. Through video analysis tracking the y-position of the tail base, we also frequently observed near-falls, in which the mouse slipped but subsequently recovered its position (Fig. 4a-b), indicating motor errors were corrected. The number of near-falls decreased over days of training, even as mice spent longer on the rotarod and higher speeds were reached (Fig. 4c). The range of 2-dimensional space taken up by movements of the tail base also decreased from Day 1-8 of training (Fig. 4d). Together, we interpreted decreases in these measures as indicators of increasingly controlled and stereotyped performance with training (i.e. fewer small motor errors).

**Fig. 4:**
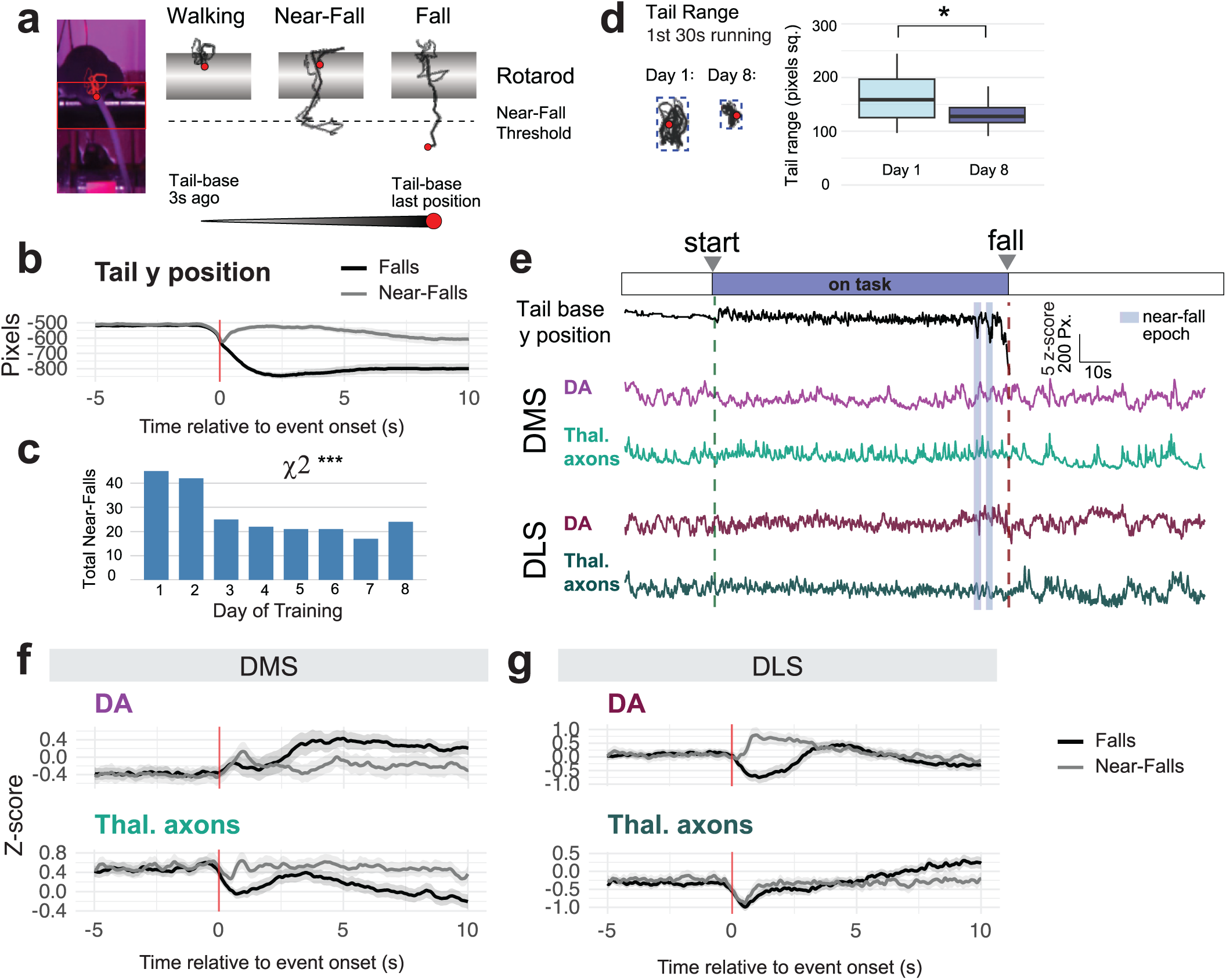
Thalamus Does Not Drive DA Error Signals in Striatum. a, Strategy for tracking tail base during accelerating rotarod, including threshold for near-falls and measurement of “Tail range.” b, Average tail base y position during falls (black line) and near-falls (gray line), n=6 mice. c, Total number of near falls per day of training, Chi square test X^2^(7) = 27.98, p<0.001***. d, Tail range measured over the first 30s of each trial lasting longer than 30s, d1 and d8 Mann-Whitney U test, W=477, p=0.023*. e, Example rotarod trial with labeled start, fall, and near-fall epochs during simultaneous position tracking and fiber photometry in DMS and DLS. f, Average photometry signals from DMS during falls (black lines) and near-falls (gray lines). g, Same as f, but in DLS.

We analyzed DA signals during falls and near-falls and found that they differed considerably across striatal subregions (Fig. 4e-g, S6). In DLS, DA signals during falls and near-falls were consistent with negative and positive prediction errors, respectively. DLS DA decreased after a fall, but on near-falls, there was a brief decrease followed by a sharp increase as position was recovered (Fig. 4g). The brief dip at near-falls diminished over training, such that overall, near-falls evoked DA increases in DLS, which appeared to move earlier in time over training (Fig. S6b). In DMS, both DA and thalamic axon responses changed over training, but they did so at different times and generally did not appear to be coactive, despite thalamic-evoked DA events being readily observable at other times within each trial (Fig. 4f, S6a-b). Thus, we concluded that the coupling between thalamic input and DMS DA does not support motor error signaling, which is more likely a function of DLS and the tail of the striatum^32^.

## Thalamus-evoked DA release is gated by task performance and DA history

Given that thalamus-evoked DA in DMS did not appear to support error signaling, we continued to search for factors regulating the strength of this relationship. Since we observed a relationship between the amplitude of thalamic axon activity and concurrent DMS DA in treadmill data (Fig. 2d-e), we examined this relationship more closely on the accelerating rotarod task. During rotarod performance, we observed increasing DA responses during larger thalamic peaks, but this relationship was not sustained during off-task recordings (Fig. 5a-b), and again no such task-dependent patterns were present in DLS (Fig. S7a). The gCaMP/rDA relationship during rotarod performance plateaued after thalamic peaks reached >4 Z-scores (Fig. 5b), which was also the regime in which we observed dependence on nAChRs (Fig. 2d). Therefore, we considered only these largest thalamic peaks >4 Z-scores for subsequent circuit-level or behavioral analyses of the factors regulating thalamus-evoked DA events.

**Fig. 5:**
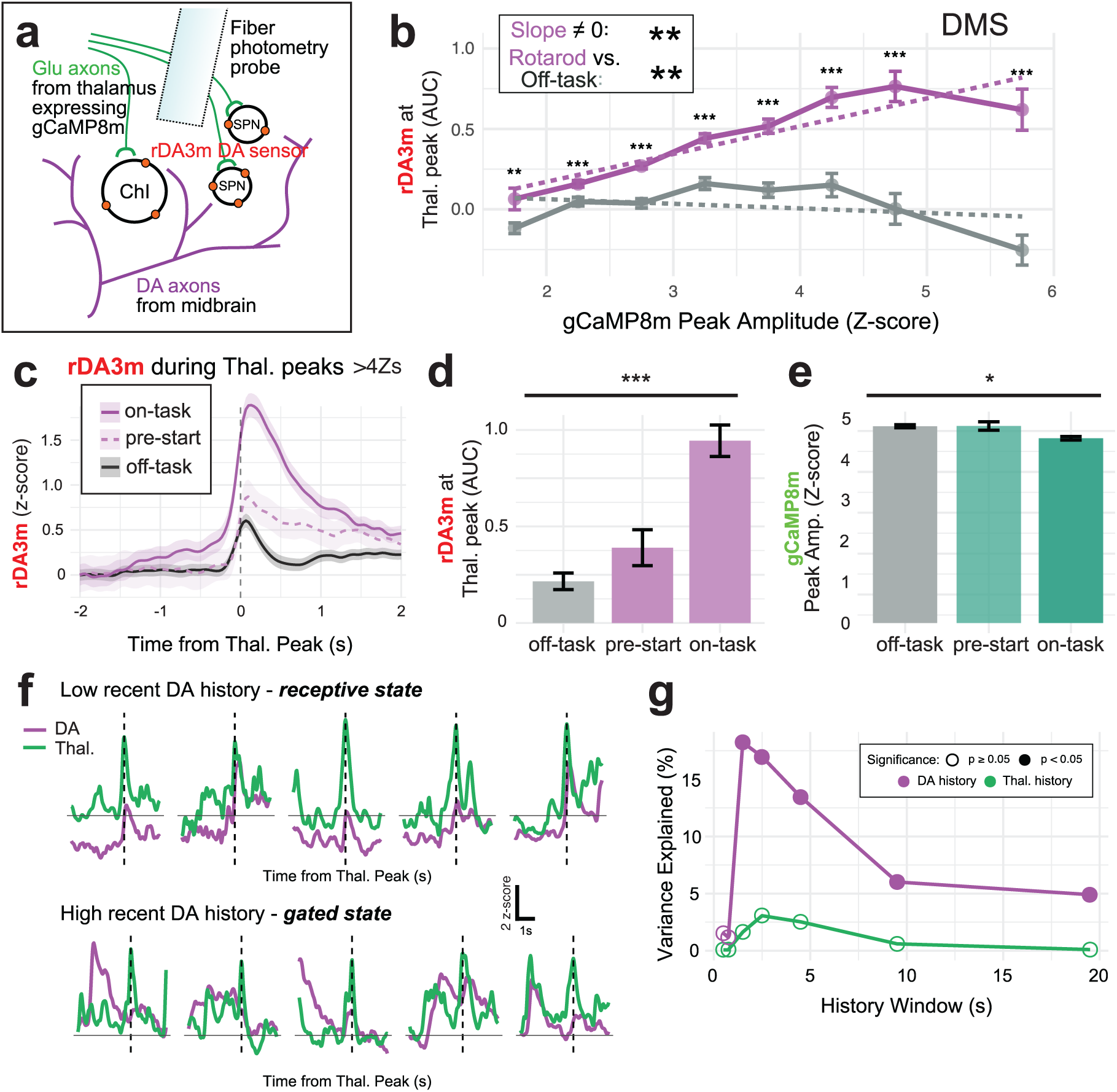
Thalamus-evoked DA release is gated by task performance and DA history. a, Diagram of sensor expression strategy for dual-color rDA3m/gCaMP8m experiments. b, Binned gCaMP8m peaks in DMS and their concurrent rDA3m responses. There is a significant effect of gCaMP8m peak amplitude on rDA3m response for rotarod trials (linear regression interaction model, see methods for details, slope=0.172, p=0.002**), but not for off-task recordings (slope= -0.029, p=0.520 ns). The relationship was significantly different in rotarod vs. off-task recordings (Fisher Z test, p= 0.005**). Bins were 0.5 Z-scores wide, and unpaired Wilcoxon Rank-Sum tests were done within each bin, with the following results for all 8 bins: p=0.007**, <0.001***, <0.001***, <0.001***, <0.001***, <0.001***, <0.001***, <0.001***). c, Traces averaging all rDA responses during gCaMP peaks >4Zs and during different behavioral epochs from 6 mice, on-task (94 peaks), the period before rod starting (71 peaks), and off-task periods (505 peaks). d, AUC quantification of c, (KW test, p <0.001***). e, Amplitudes of the same largest thalamic gCaMP8m peaks (>4 Zs) during the same task periods, showing smaller peaks during on-task periods (n as in g, KW test p=0.035*). f, Example traces of thalamic gCaMP8m (green) and rDA3m (purple) during thalamic peaks that occurred with a low recent 3s DA history value (top row) or a high recent 3s DA history value (bottom row). These traces represent peaks depicted in Fig. S7e with blue and red dots, respectively. g, Different history windows of gCaMP8m and rDA3m plotted with the variance explained of evoked rDA3m response in the large gCaMP8m peak (>4Zs) dataset. No gCaMP8m history windows explained significant variance, while all rDA3m history windows over 2s explained significant variance (linear regression, 0.5s p=0.236ns, 1s p=0.300, 2s p<0.001***, 3s p<0.001***, 5s p<0.001***, 10s p=0.017*, 20s p=0.032*).

First, we subdivided rotarod recordings into “pre-start” (when the mouse is stationary on the rotarod before trial start) and “on-task” (after rotation start) epochs. This analysis supported a strong dependence on task performance, as equivalently strong thalamic inputs evoked much larger DA responses during “on-task” vs “pre-start” epochs (Fig 5c-e).

Next, we examined several other training-related factors. We found no significant contributions of rotation speed, within-day trial number, or within-trial movement variability (calculated as the 2d bounds of tail base movements as in Fig. 4d; Fig. S4a, S7b-c).

Finally, we asked whether the recent history of the thalamic or DA signals correlated with the magnitude of thalamus-evoked DA events. We averaged the preceding Z-score values over “history windows” of different lengths (0.5s to 20s), to observe whether a given thalamus-evoked DA event was preceded by a period of low or high overall activity. The history of thalamic activity was not relevant (Fig. S7d); however, the history of recent DA activity emerged as a highly significant predictor of thalamus-evoked DA responses, with the strength of the relationship peaking at ∼3s of recent history (Fig. 5f-g). The correlation was negative (Fig. S7e), meaning that recent DA release (regardless of its coupling to thalamic input) renders the DA system relatively insensitive, or gated, to thalamic input. In contrast, if there has been relatively little recent DA release, the microcircuit is receptive to thalamic input, and a larger DA response occurs. Thus, we identified DA history as an important determinant of *in vivo* thalamus-evoked ACh-dependent DA release, which can lead to gated vs receptive states for this non-canonical mechanism of DA release.

## DISCUSSION

Since the initial demonstration that DA release in the dorsal striatum can be locally evoked via cholinergic mechanisms^8^, there has been ongoing debate about whether this pathway contributes meaningfully to striatal DA signaling *in vivo*^13,15–17,27,33^. This uncertainty is perhaps unsurprising. DA neuron firing and its downstream effects on striatal circuits form a cornerstone of computational accounts of learning and action, and an additional, locally-driven source of DA release introduces a potentially orthogonal layer of control with unknown purpose. Resolving whether and how cholinergic mechanisms engage DA release during behavior is therefore critical for understanding how dopamine supports learning, motivation, and movement.

Our findings bring clarity to this issue in four ways: (1) We show differences in the engagement of thalamus-ACh-DA mechanisms between DMS and DLS, indicating that subregional specificity is a key consideration, (2) we show that thalamic-evoked DA is dependent on behavioral context and learning, meaning that behavioral task selection and study design are key to investigating this mechanism, (3) we investigate ACh-evoked DA without relying on commonly used fluorescent ACh sensors^24^, which are less sensitive than the calcium and dopamine sensors used in our study^24–26^ and may make it difficult to resolve fine spatiotemporal ACh-DA interactions, and (4) we show that the sensitivity of the thalamus-ACh-DA circuit we studied is dependent on DA history, meaning DA release is not expected to occur at every instance of cholinergic activation, and offering insight into how separable DA release mechanisms may be coordinated *in vivo*.

### Subregional differences

Recent studies arguing against *in vivo* cholinergic stimulation of DA release or emphasizing ACh-DA anti-correlation have been collected in DLS or ventrolateral striatum^13,15,34^. Our findings are consistent with these data; we show that during motor learning and execution, thalamic inputs evoke DA release only in DMS, not in DLS. Furthermore, brief localized co-activations may often arise on a background of general anti-correlation^9^. In fact, we show that thalamic input to DLS often exhibits the reverse relationship with DA, with slight decreases in DA following thalamic transients. This depression of DLS DA release is potentially consistent with recent work showing that nAChR activation can create a refractory period in DLS dopamine terminals^34^. Indeed, sampling across sub-regions has shown that ACh-DA anti-correlation is strongest in DLS^14^, and such attention to sub-regional differences will be crucial for further elucidating the detailed mechanisms of these interactions.

### Behavioral context determines local DA release rules

NAChR activation can directly induce action potentials and evoke DA release^9^, or may render DA axons refractory to incoming input^34^. Behavioral context may be a key determinant of whether cholinergic mechanisms inhibit or promote DA release^16,17,33^. Indeed, we observed much stronger coupling between thalamic activity and DMS DA release when mice were performing the rotarod task (on-task), but not during control recordings in which they were spontaneously moving in the bottom well of the rotarod apparatus (off-task). There are several potential mechanisms. One is that changes in GABAergic tone control the gain of ACh-evoked DA, since it was recently shown that GABA_A_ receptors on DA axons gate local DA mechanisms^35^.

While there were characteristic changes to DMS and DLS DA during motor errors (near-falls and falls), these DA events were not coupled to thalamic activity, suggesting a canonical origin for DA error signals in the midbrain. This negative finding is in keeping with data from songbirds, where ACh-dependent elevations of DA during singing appeared permissive for song learning but not instructive, and DA alone reflected error signals while ACh did not^17^.

### Changes in the learning striatum

We observe that both task engagement and learning induce changes in the state of local ACh-DA interactions, altering the receptivity of DA terminals to locally-induced release. Based on our observation that thalamic-evoked DA release does not encode motor error (Fig. 4), and that thalamic activity has been related to skilled execution^36,37^, we hypothesize that locally-evoked DA release becomes more prominent with rotarod learning because motor errors decrease over training and are replaced with smooth but effortful performance. This hypothesis is supported also by recent findings suggesting a role for ACh-dependent DA release in the nucleus accumbens in promoting high-effort reward-seeking^16^, and showing that DMS dopamine released during a cholinergic burst predicts the vigor of contralateral movement^9,33^.

Future work (with consideration for subregional specificity) will be necessary to understand precisely what ACh-dependent DA release adds computationally to canonical DA signals. Although we did not observe locally induced DA release in DLS in our study, we cannot rule out the possibility that it will be observed under different behavioral conditions not yet identified. Our synaptic physiology data indicate that thalamic inputs to DLS ChIs are equally as strong as inputs to DMS ChIs, suggesting that there may be distinct rules in DLS, downstream of the thalamus-ChI synapse, that govern receptive vs gated states in this subregion.

### Technical considerations for direct measurement of ACh and DA *in vivo*

Striatal DA-ACh interactions are complex, with several distinct biological processes operating in parallel. ACh-dependent DA release has been difficult to observe during *in vivo* fiber photometry experiments employing fluorescent ACh and DA sensors together. Indeed, the dominant DA-ACh relationship observed in such experiments has been anti-correlation^13–15^, in part due to DA inhibition of ChIs via D2 receptors. However, *ex vivo* experiments have shown that cholinergic mechanisms strongly modulate DA release^38^, and can depolarize DA terminals strongly enough to initiate action potentials^9,11^. Why has it proven so difficult to detect such effects *in vivo*? Use of available fluorescent ACh sensors (e.g. GRAB-ACh3.0) in combination with fluorescent DA sensors is appealing because it allows for direct simultaneous measurement of ACh and DA *in vivo*. However, this approach has the disadvantage that GRAB-ACh3.0 is much less sensitive than the calcium or DA sensors used here. Despite roughly similar *in vivo* concentration ranges of ACh and DA in the striatum^39–42^, GRAB-ACh3.0 is ∼15x less sensitive to ACh than rDA3m is to DA (GRAB-ACh3.0 EC_50_=2000nM^24^ while GRAB-rDA3m EC_50_=130nM^25^).

Slice imaging of ACh and DA release has also revealed that spontaneous ACh events have 3-4x smaller spread than the area covered by a subsequent DA release event^9^. Therefore, a detectable incoming excitatory (e.g., thalamic) input may induce ACh release in a relatively small area of the striatum, which is difficult to detect in a bulk photometry signal transmitted by a lower-affinity sensor, as the signal is likely to be drowned out by much larger tonic ACh fluctuations. Even though undetected by fiber photometry, this limited local ACh release event could then depolarize local DA axons and evoke DA release in a significantly larger area, producing a DA signal easily detectable by currently used fluorescent DA sensors, and giving the appearance of a DA transient that is not related to any recorded ACh signal. Others have also recently argued for a temporal consideration: that *in vivo* ACh events giving rise to nAChR-dependent DA release occur too fast for most fiber photometry acquisition sampling rates to resolve the nicotinic component from a slower non-nicotinic component^27^. For these reasons, negative findings from fiber photometry experiments should be interpreted with caution, including our own negative findings on DLS. It remains possible that distinct subregional ACh-DA dynamics in DLS could obscure locally evoked DA events recorded in photometry experiments.

### Coordination of DA release mechanisms

Our data show that thalamus can drive local DA release, but thalamic input certainly does not drive all DA release. To what extent do locally-triggered and soma-triggered DA release mechanisms compete or coordinate with each other? Synthesizing our data with findings in the literature, we propose that the coordination of these mechanisms depends critically on timing. We found that thalamic-evoked DA release was effective only when there were few recent DA events (Fig. 5). This may be because ACh-DA relationship is bi-directional; low DA history may be necessary to permit strong ChI activation by relieving inhibitory D2R signaling. Indeed, ACh can either facilitate or prevent DA release depending on context^34^.

There are two possibilities for the coordination of local and soma-triggered DA signals : 1) Locally-triggered and soma-triggered DA events could represent two fully independent DA release mechanisms occurring at separate times. These orthogonal release mechanisms could correspond to computational functions in movement vigor and learning, respectively^33^. Alternatively, 2) if excitatory input to ChIs is synchronized with an incoming DA action potential, then the local cholinergic mechanism could boost soma-triggered release events. It is easy to see how additive sub-threshold depolarization could boost release. However, coordination between DA release mechanisms seems less likely if nAChRs trigger DA release by generating *de novo* action potentials at terminals^9,11^, which could collide with somatic action potentials and render DA axons refractory. Nevertheless, due to the tremendously arborized structure of DA axons (an estimated 15,000 branch points per DA neuron in mice^43^), it is possible that action potential failures at DA axon branch points regularly shape striatal patterns of DA release and that local recruitment of action potentials in additional branches has a net effect of boosting rather than canceling subsequent DA release. Additive DA release mechanisms could bring dopamine functions in movement vigor and learning together, boosting the vigor of specific valued movements or invigorating high-effort behaviors, which would be consistent with our work and others^16,33^. If future data in other behavioral contexts support a stronger mechanism for locally-evoked DA in DMS in comparison to DLS, it will be informative to scrutinize the relevant molecular and anatomical differences between subregions that create this difference.

### Other sources of ChI excitation and inhibition in vivo

In this study, we focused on thalamic inputs as a likely driver of the strong, coordinated ChI activity required to achieve ACh-dependent DA release. However, ChIs also receive other inputs. For example, a recent preprint found that frontal cortical inputs can strongly activate ChIs in DMS^44^. Stimulation of DA axons themselves can cause different burst-pause firing patterns in ChIs from different striatal subregions, at least in part due to subregional variations in glutamate co-release from DA axons^45^. In summary, although the same principles of regulation likely apply across the striatum, differing sources of ChI excitation and inhibition in striatal subregions may lead to differences in behavioral engagement of ACh-dependent DA release mechanisms across different tasks.

## Conclusion

Our findings identify thalamic input as a physiological trigger of ACh-dependent DA release in the learning striatum. We show that the engagement of this mechanism is subregion- and task-specific, and that the impact of this mechanism on overall DA signaling is dynamically gated by recent dopaminergic history. Thus, our results help clarify the mechanisms by which local and soma-driven DA signals are coordinated during behavior. By resolving whether and when cholinergic mechanisms contribute to DMS DA release *in vivo*, these results refine prevailing models of dopaminergic regulation of striatal function and suggest that DA signaling reflects an interaction between ongoing signals for vigor and learning. This framework provides a foundation for understanding how disruptions in thalamostriatal or cholinergic signaling may alter DA-dependent behaviors in neurological or psychiatric disorders.

## METHODS

### Animals

Male and female C57BL/6J (Jax #:000664), Chat-IRES-cre (Jax #:031661), or Chat-IRES-cre x Ai14 (Jax #:007914) mice were housed under a 12:12 h light/dark cycle with ad libitum access to food and water. Adult mice at least 10 weeks of age were used for all behavioral experiments, with surgeries occurring at >6 weeks of age. All experiments were approved by the Northwestern University Institutional Animal Care and Use Committee.

### Stereotactic surgeries

Viral injections and optic fiber implant surgeries were performed on mice >6 weeks of age. Mice were anesthetized in an isoflurane chamber at 3-4% isoflurane (Henry Schein) and then placed on a stereotaxic frame (Stoelting). Anesthesia was maintained at 1-3% isoflurane. Mice were injected with meloxicam (Covetrus, 20 mg/kg) subcutaneously prior to the start of surgery to minimize post-surgical pain. Hair was removed from the top of the head using Nair. The exposed skin was disinfected with alcohol and a povidone-iodine solution. Prior to incision, bupivacaine (Hospira, 2 mg/kg) was injected subcutaneously at the incision site. The scalp was opened using a sterile scalpel and holes were drilled in the skull at the appropriate stereotaxic coordinates. Viruses were infused at 50 nL/min through a blunt 33-gauge injection needle using a syringe pump (World Precision Instruments). The needle was left in place for 5 min following the end of the injection, then slowly retracted to avoid leakage up the injection tract. For photometry experiments, WT or Chat-IRES-cre mice were injected with 500nL AAV9-syn-jGCaMP8m-WPRE (2e12 vg/mL Addgene: 162375-AAV9) or AAV5-CAG-EYFP (2.5e12 vg/mL Addgene: 104055-AAV5) in thalamus at coordinates (in mm from bregma throughout): AP: -2.0, ML: 0.7, DV:-3.4. In experiments monitoring DA release in WT animals, 500nL rAAV-hSyn-rDA3m (5.5e12vg/mL, Biohippo: PT-4746) or AAV5-CaMKII-mCherry (3.3e12 vg/mL, Addgene: 114469-AAV5) was injected into DMS (0.8, 1.5, -2.8) and the same in DLS (0.3, 2.5, -3.3) before implanting fiber photometry probes. In experiments instead monitoring calcium activity in ChIs, 500nL AAV1-CAG-FLEX-NES-jRCaMP1b (2.6e12 vg/ml, Addgene: 100850-AAV1) was injected in DMS before probe placement. Fiber optic implants (Doric Lenses; 400 μm, 0.48 NA, 1.25 mm ferrules) were placed in DMS and DLS at the sites of viral injections (only in DMS for jRCaMP1b experiments). Implants were secured to the skull with Metabond (Parkell) and Flow-it ALC blue light-curing dental epoxy (Pentron). For patch clamp experiments, 500nL AAV5-CamKIIa-hChR2(H134R)-EYFP (1.2e12 vg/mL, Addgene: 26969-AAV5) was injected in thalamus (same coordinates as photometry experiments). After surgery, mice were allowed to recover until ambulatory on a heated pad, then returned to their homecage with moistened chow or DietGel available. The mice were checked after 24 hours and provided with another dose of meloxicam. All mice recovered for three to four weeks before behavioral experiments or electrophysiology recordings.

### Histology

Mice received i.p. injections of Euthasol (Virbac, 1mg/kg) to induce a rapid onset of unconsciousness and death. Once unresponsive to a firm toe pinch, an incision was made up the middle of the body cavity. An injection needle was inserted into the left ventricle of the heart, the right atrium was punctured, and solution (PBS followed by 4% PFA) was infused into the left ventricle as the mouse was exsanguinated. The mouse was then decapitated, and its brain was removed and fixed overnight at 4°C in 4% PFA. After perfusion and fixation, brains were transferred to a solution of 30% sucrose in PBS (w/v), where they were stored for at least two overnights at 4°C before making coronal sections. Tissue was sectioned on a freezing microtome (Leica) at 30 μm, stored in cryoprotectant (30% sucrose, 30% ethylene glycol, 1% polyvinyl pyrrolidone in PB) at 4°C until mounting (Fisherbrand, Cat. No. 1255005) with DAPI Fluoromont-G (Southern Biotech). Slides were imaged using a fluorescent microscope (Keyence BZ-X710) with 5x, 10x, and 40x air immersion objectives. Probe placements and injection sites were determined by comparing their location to the Allen Mouse Brain Atlas.

### Behavioral tasks and pharmacology tests

To prevent an early performance plateau during chronic accelerating rotarod training, we modified a standard mouse rotarod (Ugo Basile) to have less friction using electrical tape. To minimize movement artifacts in the photometry recordings, we balanced the patch cord on a shelf above the animal with enough slack to reach the bottom well where they would fall and collected an isosbestic control wavelength (405nm), insensitive to gCaMP and rDA, but that was used to assess and correct movement artifacts around the fall (see below). After beginning data collection, mice were placed on the stationary rod for 30s before the 4RPM to 40RPM acceleration began over the course of 300s or until the animal fell. After falling, photometry recording continued for >30s. For Mec. experiments, after 8 days of rotarod training, mice had a 4RPM treadmill test on the rotarod device, in which mice walked at a continuous slow speed for >12 minutes, which does not generally incur falls. These tests employed a Pre/Post design with a Pre recording made, followed by an I.P. injection of Mec. (Tocris, 3mg/kg) or saline vehicle, and after 20 minutes waiting, a Post recording was made also for >12 minutes. For rDA sensor block controls, SCH-23390 (Tocris, 10mg/kg) was given I.P. during a recording, then assessed post-hoc for pre-post activity.

### Behavioral Analysis

Mouse pose estimation was conducted using SLEAP. The base of the tail was manually labeled on random frames to generate initial models. Subsequently, one to two hundred labels were added at a time to generate intermediate models and assess accuracy prioritizing poorly predicted poses, and data were exported and aligned to photometry data in R. Near-falls were detected as deviations of 90 pixels in the y dimension from the average y position during the first 20s of video frames. Tail range was calculated as the diagonal length of the rectangle covered by the x/y movement of the tail base during a defined time window (e.g. during the first 30s of on-task time or the 3s preceding a photometry peak). Tail range quantification excluded any trials with falls or near-falls in the first 30s.

### Whole-cell patch clamp electrophysiology

Acute brain slices were prepared from adult mice with previous injections of AAV-ChR2-EYFP in thalamus (described above). Mice were transcardially perfused with ice-cold N-Methyl-D-Glucamine (NMDG) solution containing (in mM): 92 NMDG, 2.5 KCl, 1.2 NaH2PO4, 30 NaHCO3, 20 HEPES, 25 Glucose, 5 Na-Ascorbate, 2 Thiourea, 3 Na-Pyruvate, 10 MgSO4, 0.5 CaCl2 (Millipore Sigma). All solutions used for electrophysiology were saturated with 95%O_2_/5%CO_2_, pH was adjusted to 7.3-7.4, and osmolarity to and 300±5 mOsm. The coronally-blocked brain was glued (Loctite 454) to a specimen holder and immersed into ice-cold NMDG ACSF. Coronal slices (300 μm thick) were made using a vibratome (Leica, VT1200S). Slices were allowed to recover for 45 min in three 15 min baths: warm (33°C) NMDG solution; warm (33°C) ACSF, and lastly RT ACSF. ACSF for recovery and recordings contained in mM: 125 NaCl, 26 NaHCO3, 1.25 NaH2PO4, 2.5 KCl, 1 MgCl2, 2 CaCl2, 11 Glucose. During recordings, fresh ACSF was warmed to 30-32°C with an inline heater (Warner Instruments). To facilitate measurement of AMPA:NMDA ratios and AMPAR rectification, a cesium-based internal solution was used containing in mM: 117 CsMeSO3, 2.8 NaCl, 0.4 EGTA, 20 HEPES, 4 Mg-ATP, 0.3 Na3-GTP, 5 QX314 Cl, 5 TEA-Cl, 0.1 Spermine 4HCl. The following drugs were added to the recording ACSF: picrotoxin (50 μM, Sigma), and to measure AMPA rectification indices, D-AP5 (50 μM, Tocris). Patch pipettes (3-5 MΩ) were pulled (Narishige, PC-100) from borosilicate glass (Warner Instruments, G150TF-4) and moved with the assistance of a micromanipulator (Sensapex). Cells were visualized with a 40x water-immersion objective (NA 0.8, Olympus, #N2667700) on a microscope (Olympus, BX51WI) equipped with infrared-differential interference imaging (DIC) and a camera (QImaging, Retiga Electro Monochrome). An LED light source (CoolLED, pE-300^white^) was used to illuminate the slice through the objective for targeted patching and for 1ms optogenetic stimulation. With the aid of a power meter (Thor Labs, PM130D), the LED power was adjusted to deliver ∼5mW/mm² at 475 nm to the slice. Signals were recorded at 10 kHz using Wavesurfer v0.945 (https://wavesurfer.janelia.org/), a National Instruments Digitizer (NIDAQ X series PCIe-6323) and BNC Breakout (BNC-2090A), and a Multiclamp 700B amplifier (Molecular Devices). All experiments were done in voltage clamp mode, holding at -70mV or +40mV for AMPA:NMDA ratios and rectification indices. Data analysis was performed offline using custom-written R scripts.

### In Vivo Fiber Photometry

Photometry data were collected and synchronized with behavioral data using Tucker Davis Technologies (TDT) Synapse with iCon v2. All recordings were performed using a fiber photometry rig with optical components from Doric lenses and TDT controlled by a RZ10X real-time processor. 405nm, 465nm, and 560nm LEDs were modulated at 210 Hz, 330 Hz, and 450 Hz, respectively. LED currents were adjusted to return a voltage between 150-200mV for each signal, were offset by 5 mA, were demodulated using a 4 Hz lowpass frequency filter. GuPPy, an open source Python-based photometry data analysis pipeline, was used to process fiber photometry data into .csv format. The 405nm isosbestic (signal insensitive) channel was monitored to evaluate fluctuations due to movement artifacts, especially around the falls on the rotarod, but these were minimal. Nevertheless, for analysis of photometry data during falls and near-falls, isosbestic subtraction was performed in GuPPy. To facilitate comparisons across animals, z-scores were calculated by subtracting the mean ΔF/F calculated across the entire session and dividing by the standard deviation (GuPPy standard z-score method). Signals were then averaged around the nearest video frame, downsampling to the video frame rate of 30Hz for further analysis.

### Quantification and Data Analysis

All subsequent data analyses were performed in R (v. 4.5.1) and visualized using ggplot2. Z-scored peaks were identified and quantified using the R pracma package (findpeaks) with minimal inter-peak separation of 2s. Area under the curve (AUC) was calculated in R using trapezoidal integration for the z-scored rDA signal using a baseline period of -2 to -0.5s relative to peak subtracted from a response window of -0.25 to +0.5s. For initial analysis of peaks, to minimize bias due to biological changes across learning and between task epochs, rather than thresholding peaks at a particular Z score value, the largest peaks were taken from each recording using roughly 5 peaks/min (giving 10 peaks per rotarod trial and 60 peaks per 12 min. treadmill recording). This method provided the whole distribution of peaks down to ∼1.5 Zs, but did not obscure changes across recordings. For correlation and history analysis of only the largest gCaMP peaks (Fig. 5), peaks were thresholded above 4Zs. For binning analyses, identified gCaMP peaks were pooled across animals and binned by gCaMP amplitude into 8 bins using predefined edges: 1.5, 2.0, 2.5, 3.0, 3.5, 4.0, 4.5, >5.0. In R, linear regression slope and Pearson R were calculated using cor.test with 6 degrees of freedom, and Fisher Z test compared correlations between groups. Bin-wise tests were unpaired Mann-Whitney U tests. For changes over days, each animal contributed 8 trials per day per condition (rotarod vs. off-task), and recording (trial)-level values were used for linear mixed-effects modeling (LMM) approaches implemented using the lme4 package (v1.1-35.1) for linear mixed model fitting and lmerTest (v3.1-3) for p-value calculation. The full model was specified as: Response ∼ as.factor(day) + trial_type + (1|animal) + (1|animal:day), where Response represents the measured variable, day is a categorical factor representing training day (1-8), and trial_type is a categorical factor with two levels (rodtrials vs. off-task). We conducted three statistical tests for each brain region (DMS, DLS) and each measured variable: 1. Effect of day on rotarod trials and 2. Effect of day on off-task trials were assessed using models made from only the relevant trial type: Response ∼ as.factor(day) + (1|animal) + (1|animal:day), with Type II ANOVA F-test on the day coefficient, while 3. Overall difference between Rotarod and off-task trials employed the full model with both conditions: Response ∼ as.factor(day) + trial_type + (1|animal) + (1|animal:day), with Type II ANOVA F-test on the trial_type coefficient. ANOVA was implemented via the anova() function in lmerTest, which provides F-statistics and p-values using Satterthwaite’s method for approximating degrees of freedom. For assessing variance explained by gCaMP or rDA history, mean Z-score value was calculated in the specified time window and compared to the DA response AUC via linear regression: DA response AUC ∼ signal history, from which we extracted R² and tested significance using F-statistic from the linear model, F = (MS_regression) / (MS_residual). For whole-cell patch clamp data, AMPA:NMDA ratios were calculated using the EPSC amplitude at -70mV and the response at +40mV at 50ms after stimulation. AMPA rectification indices were calculated using responses at -70mV divided by responses at +40mV in the presence of 50μM D-AP5.

The data, code, protocols, and key lab materials used and generated in this study are listed in the Key Resource Table alongside their persistent identifiers.

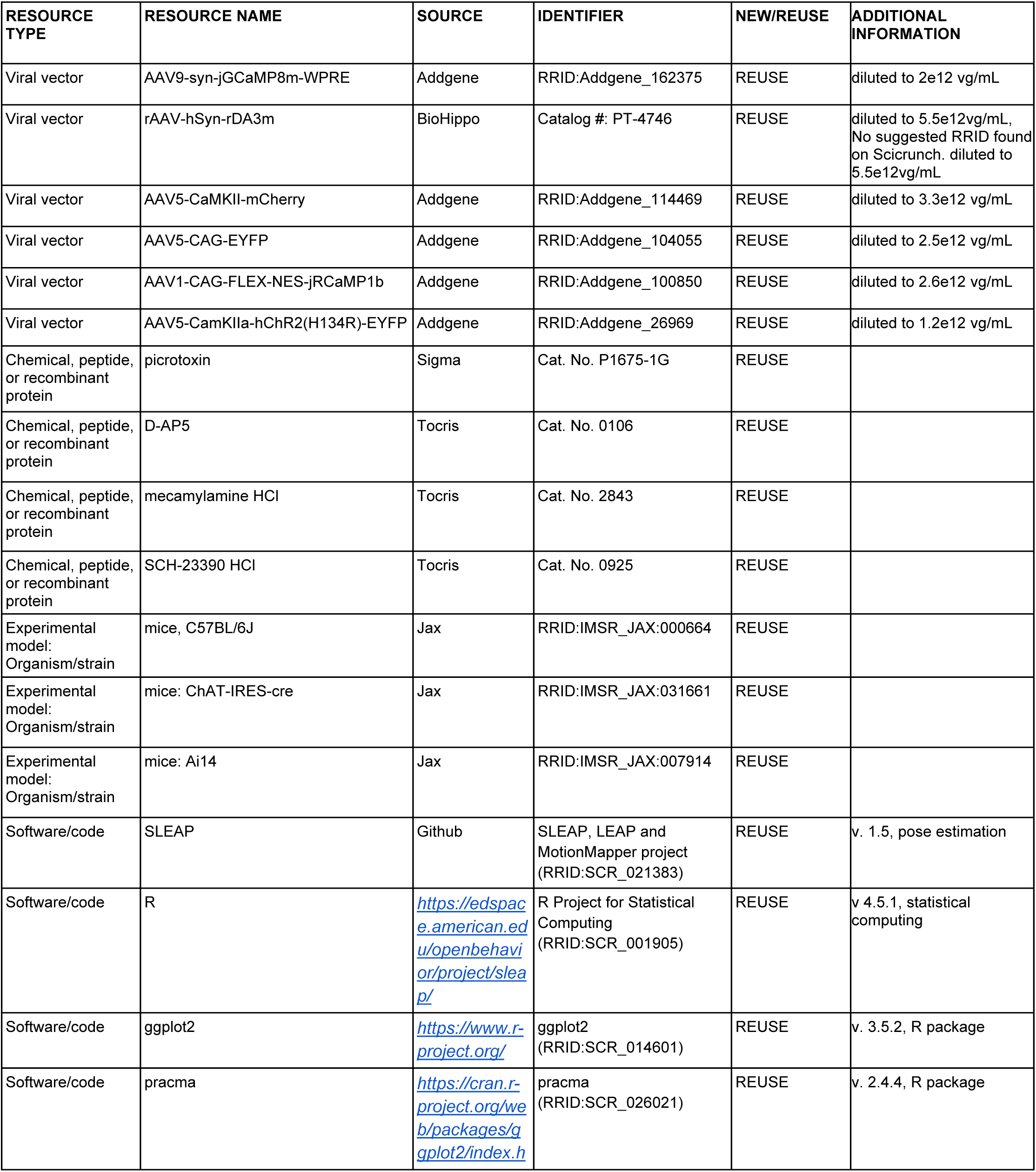
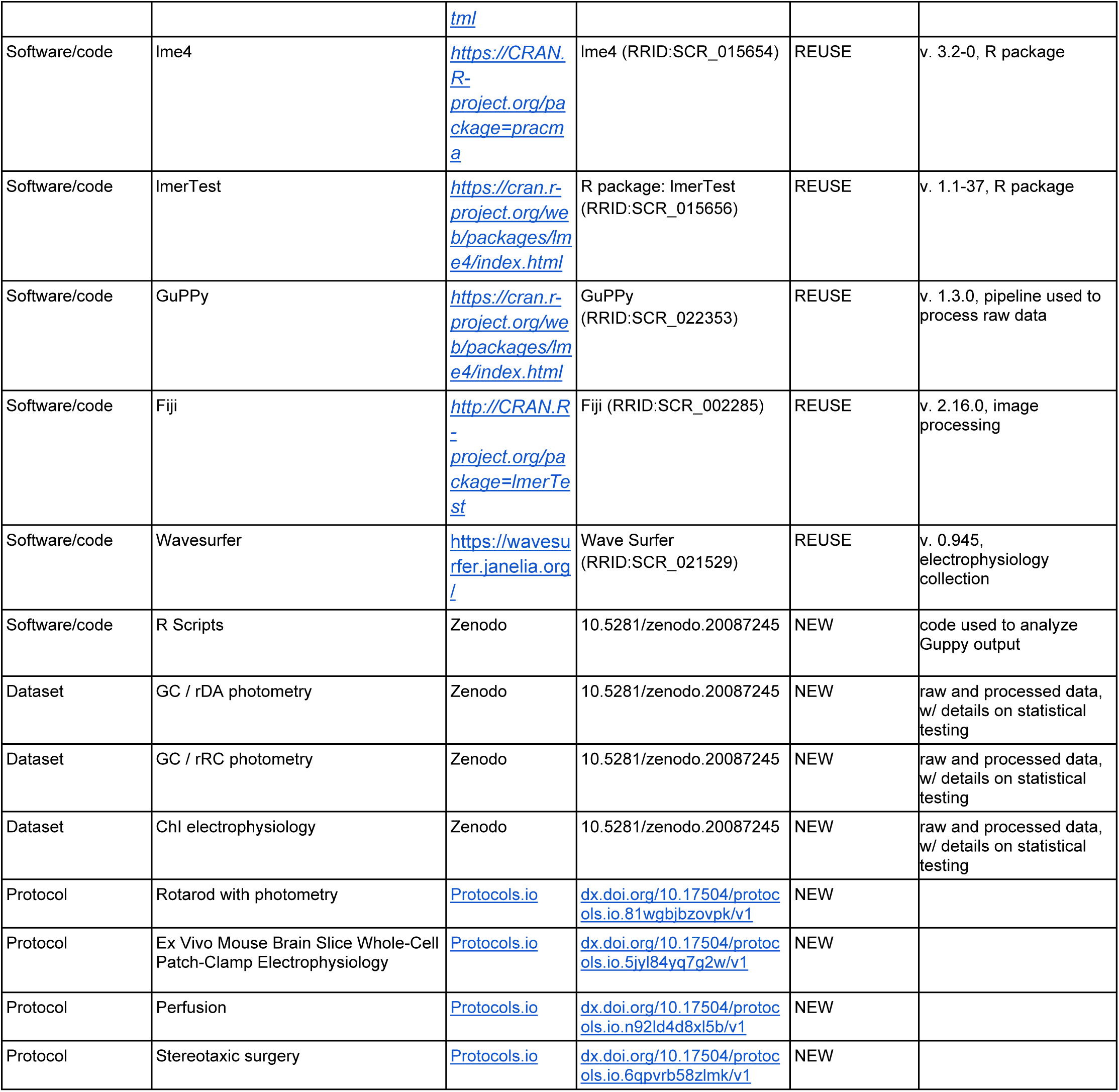
KEY RESOURCE TABLE.

## ACKNOWLEDGEMENTS

We thank the Lerner laboratory and members of “Team Edwards” from the Aligning Science Across Parkinson’s (ASAP) Collaborative Research Network for helpful discussions and critical feedback throughout the project. We thank Louis Van Camp for supporting the laboratory’s general operations throughout the project. We thank Venus Sherathiya for technical support using GuPPy for fiber photometry analysis. We thank the Center for Comparative Medicine at Northwestern University for providing animal care and husbandry.

## AUTHOR CONTRIBUTIONS

A.J.M. and T.N.L. conceived the studies and wrote the manuscript. A.J.M., M.Z., and B.D. conducted the experiments. A.J.M. and R.F.K. analyzed the data. T.N.L. provided oversight and support for the project.

## MATERIALS & CORRESPONDENCE

Correspondence and requests for materials should be addressed to Talia N. Lerner.

## SUPPLEMENTAL FIGURE LEGENDS

**Fig. S1:**
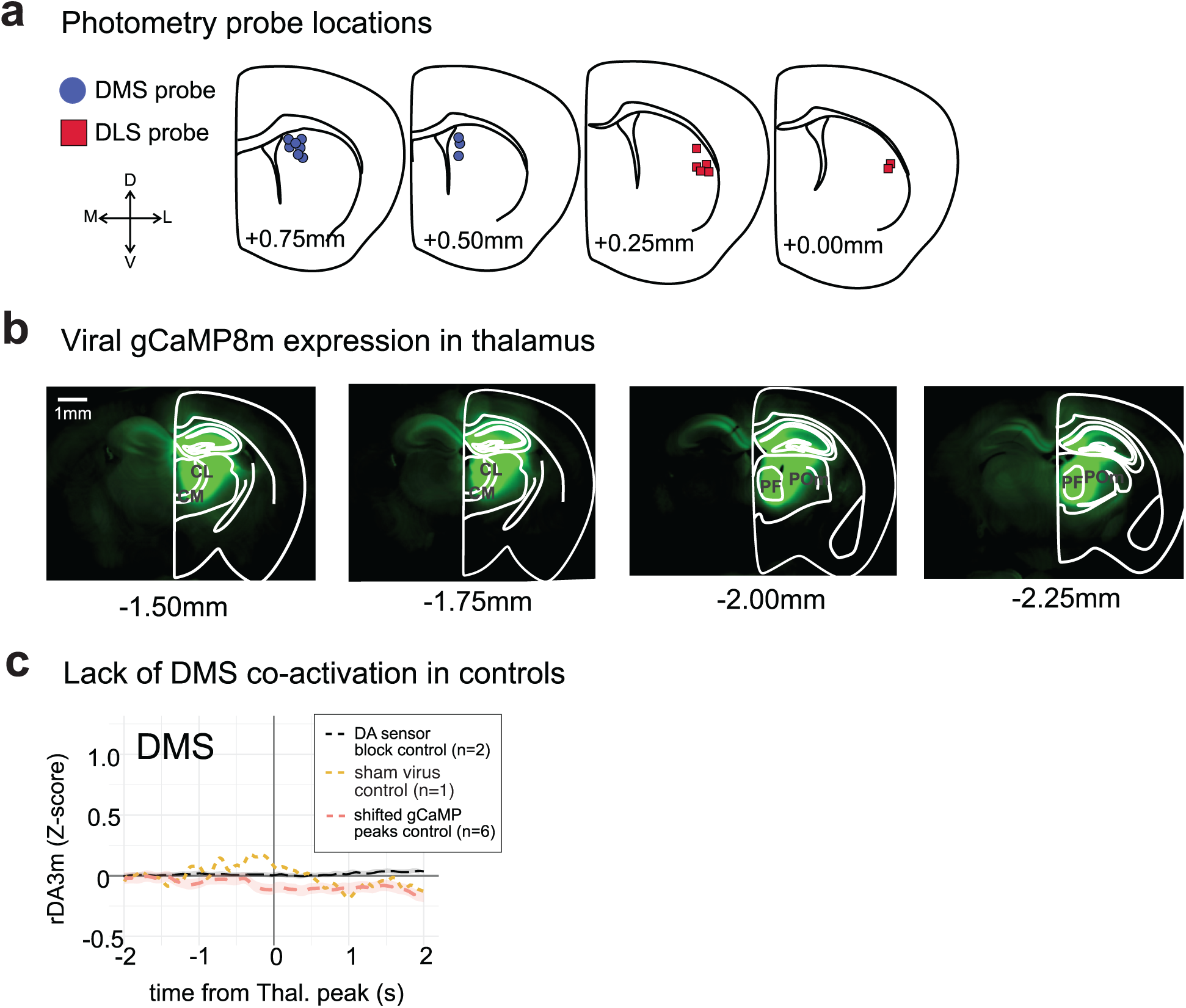
Anatomical and control information for photometry experiments in DMS and DLS. a, DMS and DLS probe placements for all animals included in photometry experiments, snapped to nearest 250 μm plane. b, Example thalamic gCaMP8m injection showing rostrocaudal detail. Thalamic injections centered on the parafasicular (PF) nucleus, but also included rostral intralaminar central median (CM) and central lateral (CL) nuclei, which also project to striatum and were not considered separately in this study. c, Controls for dual-color photometry in DMS, signal processing, and data analysis pipeline. Black dashed line is rDA3m response to thalamic gCaMP8m peaks during blockade of rDA3m’s DA binding site with SCH-23390 (10mg/kg), as in Fig. 1e. Yellow dashed line is 560nm response to “peaks” in an animal where EYFP and mCherry were expressed in place of gCaMP8m and rDA3m. Orange dashed line is rDA3m response in real trial data, but using timestamps for gCaMP8m peaks which are arbitrarily shifted forward 4s; compare response against rDA3m response during genuine gCaMP8m peak timestamps in Fig. 1f.

**Fig. S2:**
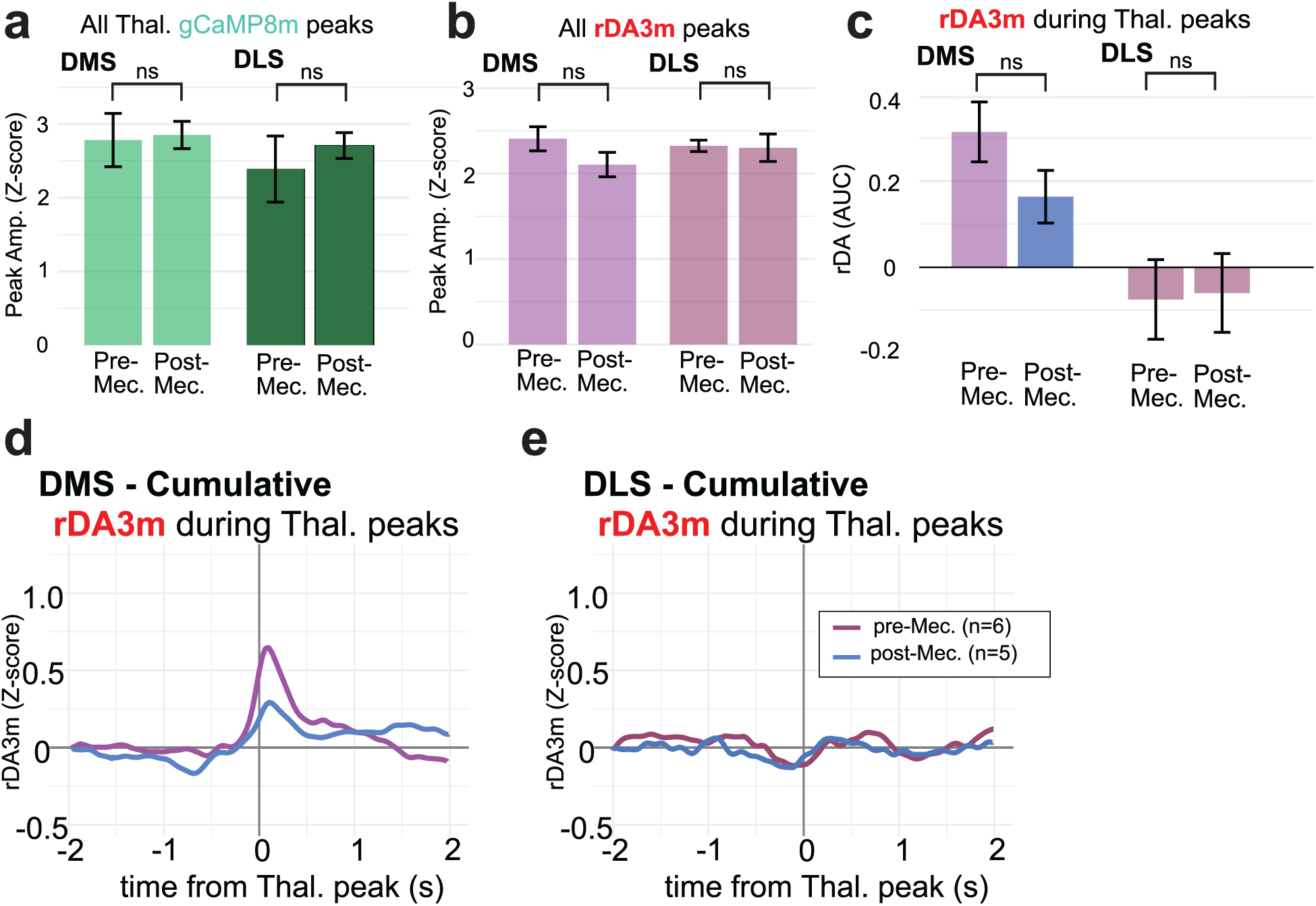
Supplemental pharmacological data in DMS and DLS. a, Thalamic axon gCaMP8m peak amplitudes before and after Mec. in DMS (paired Wilcoxon signed-rank test, p=1ns) and DLS (p=1ns) (n=5 mice). b, rDA3m peak amplitudes before and after Mec. in DMS (p=0.106ns) and DLS (p=0.787ns). c, rDA3m AUC during gCaMP8m peaks before and after MEC in DMS (p=0.106ns) and DLS (p=0.590ns). d, Direct comparison of average rDA response during gCaMP8m peaks pre- (purple) and post-Mec. (blue) in DMS, same data as in Fig 2b-c. e, Same as d, but in DLS.

**Fig. S3:**
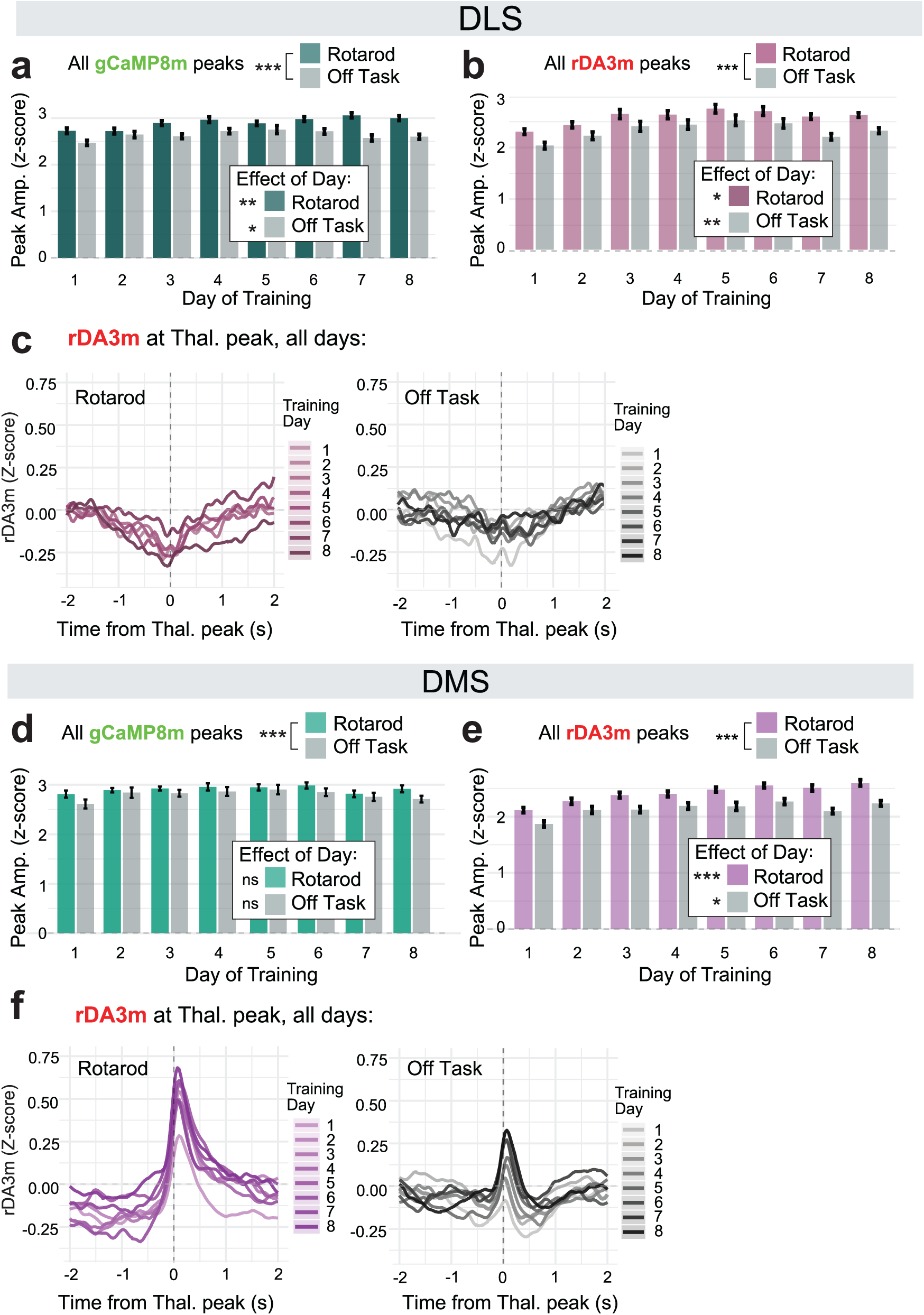
Supplemental data for changes over days of rotarod training in DLS and DMS. a, Average amplitude of thalamic axon gCaMP8m peaks in DLS during rotarod trials (green), off-task recordings (gray). Effect of training day on gCaMP8m peaks during rotarod trials (F(7, 35.03) = 3.54, p = 0.006**) and off-task recordings (F(7, 35.65) = 2.70, p = 0.024*). b, Same as a, but average rDA3m peak amplitudes in DLS. Effect of training day on rDA3m peaks during rotarod trials (F(7, 35.06) = 2.98, p = 0.015*), or off-task recordings (F(7, 34.88) = 3.50, p = 0.006**). c, rDA3m responses in DLS during thalamic gCaMP8m peaks over days of training, average from all 6 mice. Purple traces (left) for rotarod trials and gray traces (right) for off-task recordings. d, Same as a, but for DMS. GCaMP8m peak amplitudes did not change significantly across training days on rotarod (LMM, see methods for details: F(7, 34.95) = 0.88, p = 0.532 ns) or in off-task control recordings (F(7, 35.22) = 0.80, p = 0.594 ns). Rotarod recordings showed a slight but consistent trend toward larger thalamic gCaMP8m amplitudes than off-task recordings (F(1, 709.50) = 12.16, p < 0.001***; n=6 mice). e, Same as b, but for DMS. rDA3m peak amplitude increased across training days in rotarod trials (F(7, 34.96) = 5.08, p < 0.001***) and also increased in off-task recordings (F(7, 35.34) = 3.01, p = 0.014*). Rotarod trials showed larger amplitudes than off-task recordings overall (F(1, 709.29) = 137.16, p < 0.001***; n=6 mice). f, rDA3m responses in DMS during thalamic gCaMP8m peaks over days of training, average from all 6 mice. Purple traces (left) for rotarod trials and gray traces (right) for off-task recordings.

**Fig. S4:**
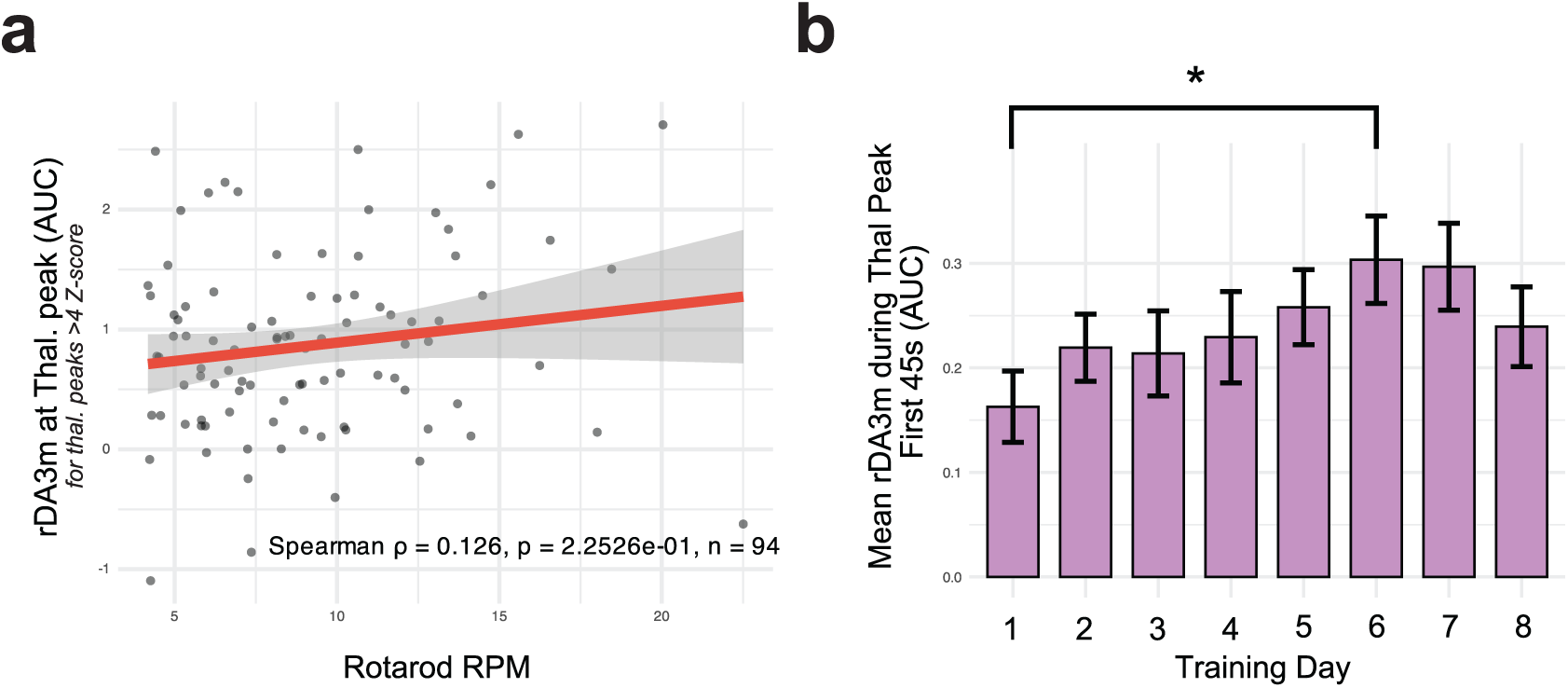
Thalamus-associated DA responses increase over days independent of running speed. a, As below in Fig. S7, data from 6 mice were restricted to analysis of >4Zs gCaMP8m peaks and concurrent rDA3m response magnitudes were analyzed. The DA response did not correlate significantly with RPM of the rotarod (n=94 peaks, Spearman ρ = 0.126, p=0.225ns). b, Because the 4-40RPM rotarod acceleration program is the same during trials on every day, the first 45s on day 1 is at the same relatively low speed as on subsequent days. When subsetting to data collected only in the first 45s of a rotarod trial, and excluding trials with falls during the first 45s, rDA3m during thalamic gCaMP8m peaks increased from day 1 and peaked on day 6, as in the non-subsetted data (c.f. Fig. 3d-e), suggesting that this increase is not due to faster running speed.

**Fig. S5:**
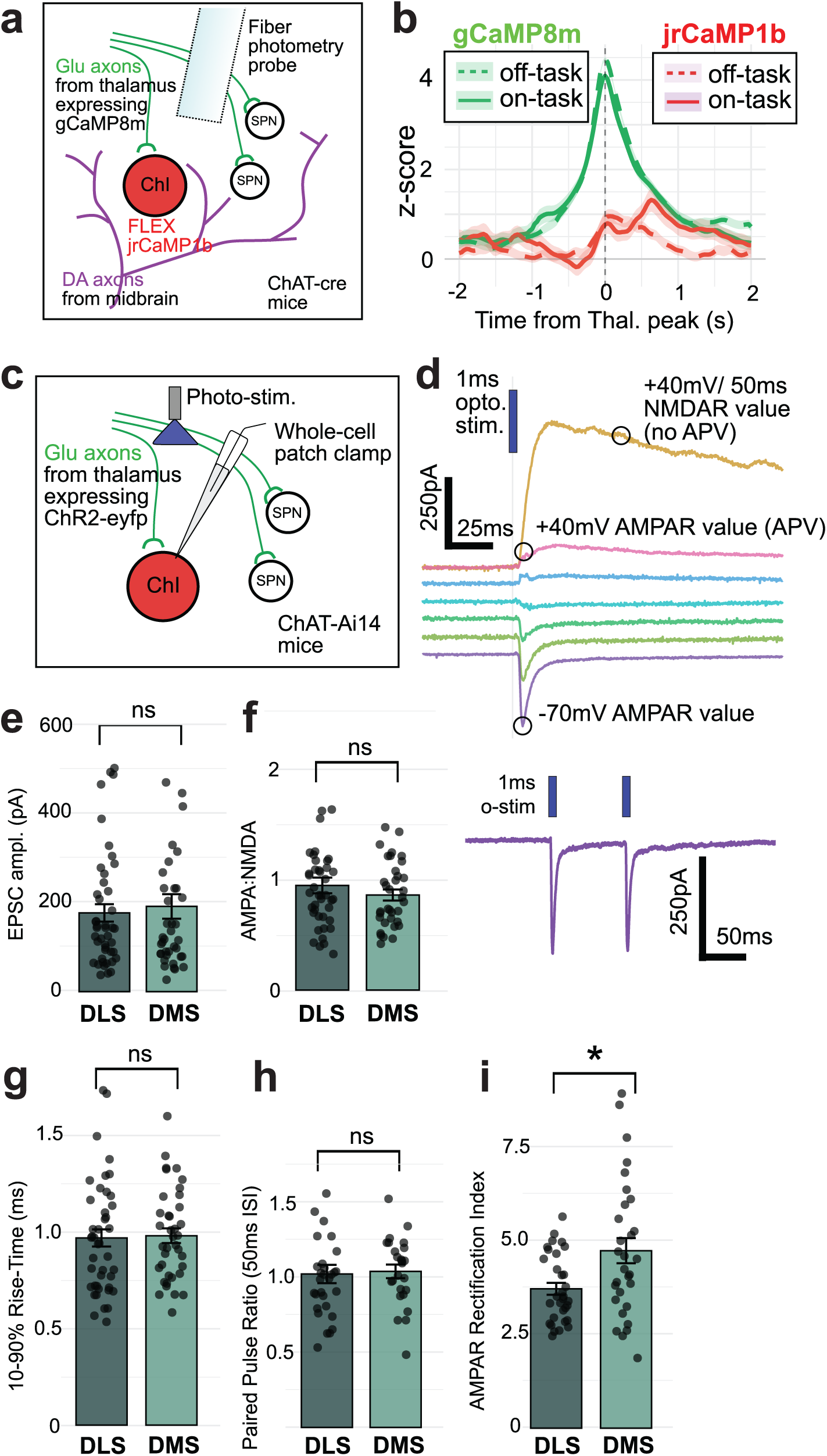
Thalamic activation of ChIs is not task-dependent, and differences in the properties of thalamostriatal synapses onto ChIs in DMS and DLS do not explain the subregional difference in thalamic-evoked DA release. a, Diagram of sensor expression strategy for dual-color jrCaMP1b/gCaMP8m experiments in DMS only. b, Thalamus-associated ChI activation is not task dependent, in contrast to the task-dependence of thalamus-associated rDA3m events (c.f. Fig. 5c). c, Experimental schematic for *ex vivo* whole cell patch clamp experiments. Patch clamp data from 84 ChIs in 22 animals. d, Example traces from a whole cell recording targeting a ChI in DLS while optogenetically activating thalamic axons to measure EPSC amplitude, rise time, AMPA:NMDA ratio, AMPAR rectification, and Paired-pulse ratio (PPR). e, EPSC amplitude (n=39 cells in DMS, 44 in DLS, Wilcoxon rank-sum test, p=0.898ns). f, AMPA:NMDA ratio (n=35,42, p=0.471ns). g, 10-90% rise time (n=38,43, p=0.623ns). h, PPR 50ms interstimulus interval (n=25,30, p=0.403ns). i, AMPAR rectification index (n=30,32,p=0.027*).

**Fig. S6:**
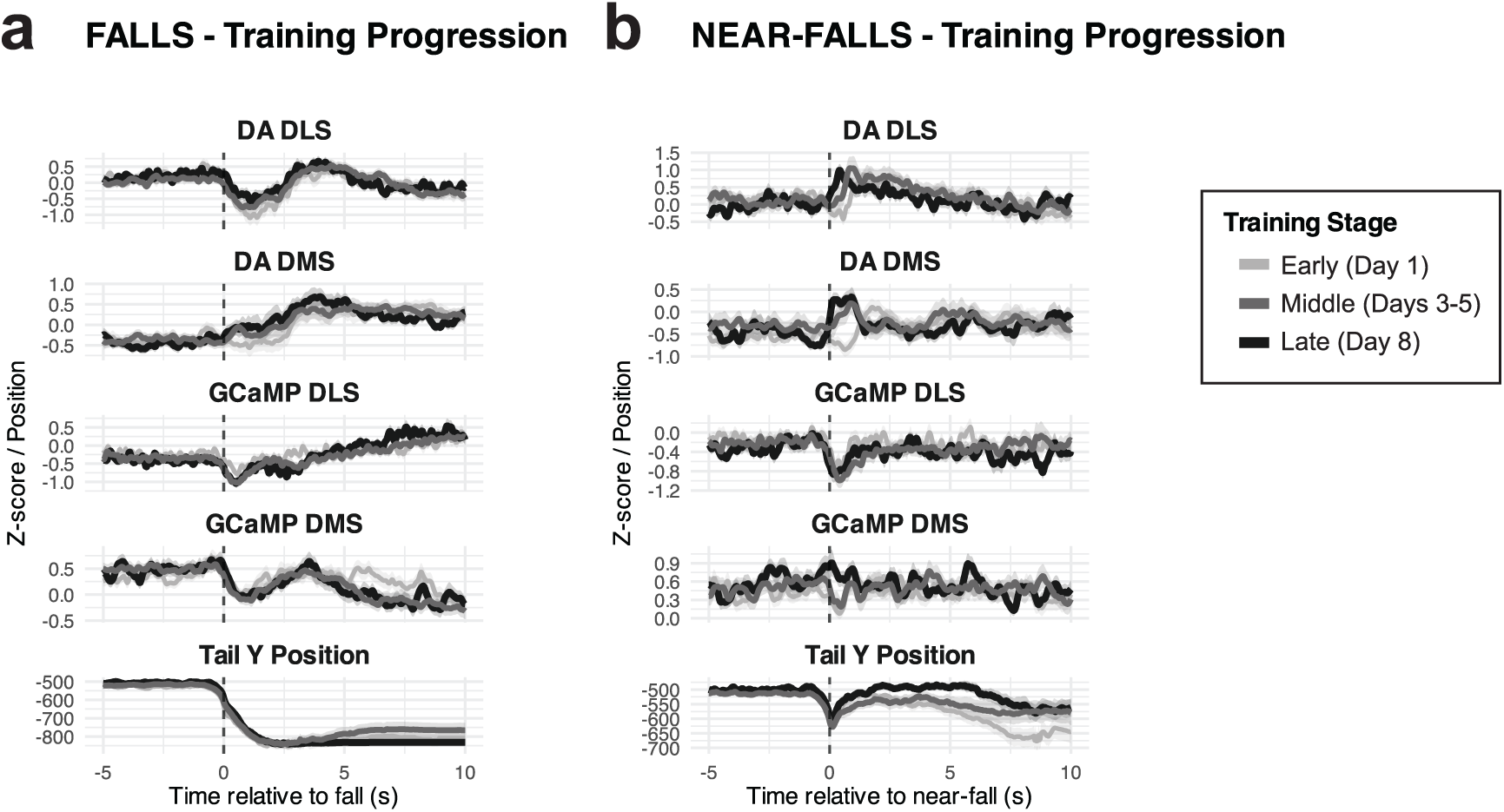
Progression of photometry signals during falls and near-falls over days of training. a, Averaged rDA3m signals or thalamic gCaMP8m signals during falls during rotarod training with y position of tail base at the bottom, separated by early training (day 1, light gray), middle training (days 3-5, gray), or late training (day 8, black). b, Same as a, but for near-falls.

**Fig. S7:**
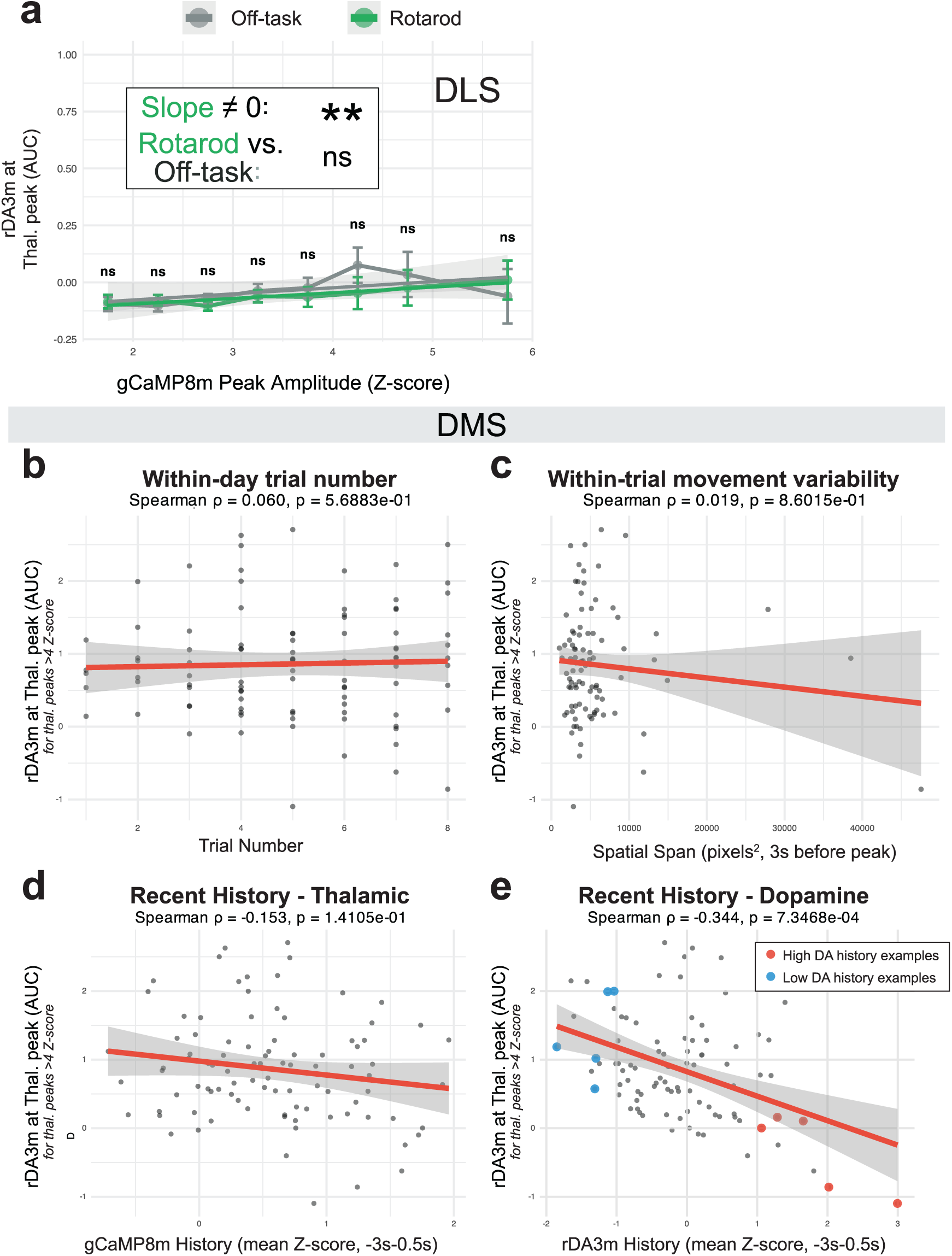
DLS does not show thalamus-evoked DA regardless of gCaMP8m amplitude or task state, and supplemental correlations with thalamus-evoked DA in DMS. a, Binned gCaMP8m peaks in DLS and their concurrent rDA3m responses (c.f. DMS data in Fig. 5b at the same scale). There is a significant linear relationship between gCaMP8m peak amplitude on rDA3m response for rotarod trials (linear regression interaction model, see methods for details, p=0.002**), but not for off-task recordings (p=0.139ns). The relationship was not significantly different in rotarod vs. off-task recordings (Fisher Z test, p= 0.157ns). Unpaired Wilcoxon Rank-Sum tests were done within each bin, all p>0.05ns. b-e, Data from 6 mice was subsetted to the largest >4Zs gCaMP peaks and concurrent rDA3m responses for analyses of determinants of rDA3m response magnitude. There was no significant correlation between rDA3m response magnitude during a large thalamic gCaMP8m peak and within-day trial number (Spearman ρ = 0.060, p=0.569ns), within-trial movement variability (i.e. tail range, Spearman ρ = 0.019, p=0.860ns), or the recent (3s) gCaMP8m history (Spearman ρ = -0.153, p=0.141ns). There was significant negative correlation between rDA3m response magnitude and recent (3s) DA history (Spearman ρ = -0.344, p<0.001***). Blue and red dots indicate rDA3m responses to large gCaMP8m peaks preceded by low and high DA history, respectively, shown as example traces in Fig. 5f.

## REFERENCES

1. Cox, J. & Witten, I. B. Striatal circuits for reward learning and decision-making. Nat Rev Neurosci 20, 482–494 (2019).

2. Wise, R. A. Dopamine, learning and motivation. Nat Rev Neurosci 5, 483–494 (2004).

3. Todd, K. L., Cramb, K. M. L., Brimblecombe, K. R. & Cragg, S. J. New insights into axonal regulators of dopamine transmission in health and disease. Current Opinion in Neurobiology 94, 103093 (2025).

4. Mirenowicz, J. & Schultz, W. Preferential activation of midbrain dopamine neurons by appetitive rather than aversive stimuli. Nature 379, 449–451 (1996).

5. Schultz, W. Predictive Reward Signal of Dopamine Neurons. Journal of Neurophysiology 80, 1–27 (1998).

6. Schultz, W., Dayan, P. & Montague, P. R. A Neural Substrate of Prediction and Reward. Science 275, 1593–1599 (1997).

7. Bayer, H. M. & Glimcher, P. W. Midbrain Dopamine Neurons Encode a Quantitative Reward Prediction Error Signal. Neuron 47, 129–141 (2005).

8. Threlfell, S. et al. Striatal Dopamine Release Is Triggered by Synchronized Activity in Cholinergic Interneurons. Neuron 75, 58–64 (2012).

9. Liu, C. et al. An action potential initiation mechanism in distal axons for the control of dopamine release. Science 375, 1378–1385 (2022).

10. Cover, K. K. et al. Activation of the Rostral Intralaminar Thalamus Drives Reinforcement through Striatal Dopamine Release. Cell Reports 26, 1389–1398.e3 (2019).

11. Kramer, P. F. et al. Synaptic-like axo-axonal transmission from striatal cholinergic interneurons onto dopaminergic fibers. Neuron 110, 2949–2960.e4 (2022).

12. Azcorra, M. et al. Unique functional responses differentially map onto genetic subtypes of dopamine neurons. Nat Neurosci 26, 1762–1774 (2023).

13. Krok, A. C. et al. Intrinsic dopamine and acetylcholine dynamics in the striatum of mice. Nature 621, 543–549 (2023).

14. Bouabid, S. et al. An anatomical hotspot for striatal dopamine-acetylcholine interactions during reward and movement. 2026.01.20.700614 Preprint at 10.64898/2026.01.20.700614 (2026).

15. Chantranupong, L. et al. Dopamine and glutamate regulate striatal acetylcholine in decision-making. Nature 621, 577–585 (2023).

16. Touponse, G. C. et al. Cholinergic modulation of dopamine release drives effortful behaviour. Nature 1–10 (2026) doi:10.1038/s41586-025-10046-6.

17. Qi, J., Schreiner, D. C., Martinez, M., Pearson, J. & Mooney, R. Dual neuromodulatory dynamics underlie birdsong learning. Nature 641, 690–698 (2025).

18. Yin, H. H. et al. Dynamic reorganization of striatal circuits during the acquisition and consolidation of a skill. Nat Neurosci 12, 333–341 (2009).

19. Dang, M. T. et al. Disrupted motor learning and long-term synaptic plasticity in mice lacking NMDAR1 in the striatum. Proceedings of the National Academy of Sciences 103, 15254–15259 (2006).

20. Lapper, S. R. & Bolam, J. P. Input from the frontal cortex and the parafascicular nucleus to cholinergic interneurons in the dorsal striatum of the rat. Neuroscience 51, 533–545 (1992).

21. Ding, J. B., Guzman, J. N., Peterson, J. D., Goldberg, J. A. & Surmeier, D. J. Thalamic Gating of Corticostriatal Signaling by Cholinergic Interneurons. Neuron 67, 294–307 (2010).

22. Mamaligas, A. A., Barcomb, K. & Ford, C. P. Cholinergic Transmission at Muscarinic Synapses in the Striatum Is Driven Equally by Cortical and Thalamic Inputs. Cell Reports 28, 1003–1014.e3 (2019).

23. Johansson, Y. & Silberberg, G. The Functional Organization of Cortical and Thalamic Inputs onto Five Types of Striatal Neurons Is Determined by Source and Target Cell Identities. Cell Reports 30, 1178–1194.e3 (2020).

24. Jing, M. et al. An optimized acetylcholine sensor for monitoring in vivo cholinergic activity. Nat Methods 17, 1139–1146 (2020).

25. Zhuo, Y. et al. Improved green and red GRAB sensors for monitoring dopaminergic activity in vivo. Nat Methods 21, 680–691 (2024).

26. Zhang, Y. et al. Fast and sensitive GCaMP calcium indicators for imaging neural populations. Nature 615, 884–891 (2023).

27. Flink, D. R., Faturos, N. G., Zhang, H. & Hamid, A. A. Dual cholinergic mechanisms for sculpting striatal dopamine in vivo. 2025.12.19.695021 Preprint at 10.64898/2025.12.19.695021 (2025).

28. Jones, E. G. The Thalamus. (Cambridge University Press, Cambridge ; New York, 2007).

29. Kaneko, S. et al. Systemic injection of nicotinic acetylcholine receptor antagonist mecamylamine affects licking, eyelid size, and locomotor and autonomic activities but not temporal prediction in male mice. Mol Brain 15, 77 (2022).

30. Götz, T. et al. Functional Properties of AMPA and NMDA Receptors Expressed in Identified Types of Basal Ganglia Neurons. J. Neurosci. 17, 204–215 (1997).

31. Suzuki, T., Miura, M., Nishimura, K. & Aosaki, T. Dopamine-Dependent Synaptic Plasticity in the Striatal Cholinergic Interneurons. J. Neurosci. 21, 6492–6501 (2001).

32. Greenstreet, F. et al. Dopaminergic action prediction errors serve as a value-free teaching signal. Nature 643, 1333–1342 (2025).

33. Jang, H. J., McMahon Ward, R., Golden, C. E. M. & Constantinople, C. M. Acetylcholine demixes heterogeneous dopamine signals for learning and moving. Nat Neurosci 29, 840–850 (2026).

34. Zhang, Y.-F. et al. An axonal brake on striatal dopamine output by cholinergic interneurons. Nat Neurosci 28, 783–794 (2025).

35. Brill-Weil, S. G., et al. Presynaptic GABAA receptors control integration of nicotinic input onto dopaminergic axons in the striatum. Cell Reports 44, (2025).

36. Wolff, S. B. E., Ko, R. & Ölveczky, B. P. Distinct roles for motor cortical and thalamic inputs to striatum during motor skill learning and execution. Science Advances 8, eabk0231 (2022).

37. Gjoni, E. et al. Complementary cortical and thalamic contributions to cell type–specific striatal activity dynamics during movement. Science Advances 12, eaea3935 (2026).

38. Rice, M. E. & Cragg, S. J. Nicotine amplifies reward-related dopamine signals in striatum. Nat Neurosci 7, 583–584 (2004).

39. Parsons, L. H. & Justice, J. B. Extracellular concentration and in vivo recovery of dopamine in the nucleus accumbens using microdialysis. J Neurochem 58, 212–218 (1992).

40. Chen, K. C. Evidence on extracellular dopamine level in rat striatum: implications for the validity of quantitative microdialysis. J Neurochem 92, 46–58 (2005).

41. Vinson, P. N. & Justice, J. B. Effect of neostigmine on concentration and extraction fraction of acetylcholine using quantitative microdialysis. Journal of Neuroscience Methods 73, 61–67 (1997).

42. DeBoer, P. & Abercrombie, E. D. Physiological release of striatal acetylcholine in vivo: modulation by D1 and D2 dopamine receptor subtypes. The Journal of Pharmacology and Experimental Therapeutics 277, 775–783 (1996).

43. Seddon, J. & Kramer, P. F. Intrinsic and synaptic regulation of axonal excitability in dopaminergic neurons. Front. Cell. Neurosci. 19, (2025).

44. Goldbach, H. C. et al. Cholinergic-dependent dopamine signals in mouse dorsal striatum are regulated by frontal but not sensory cortices. 2025.09.30.679538 Preprint at 10.1101/2025.09.30.679538 (2025).

45. Chuhma, N., Mingote, S., Moore, H. & Rayport, S. Dopamine Neurons Control Striatal Cholinergic Neurons via Regionally Heterogeneous Dopamine and Glutamate Signaling. Neuron 81, 901–912 (2014).

