## Supplemental Figures for "Thalamus orchestrates local acetylcholine-dependent dopamine release in the learning striatum"

Supplementary Figure 1

**a** Photometry probe locations

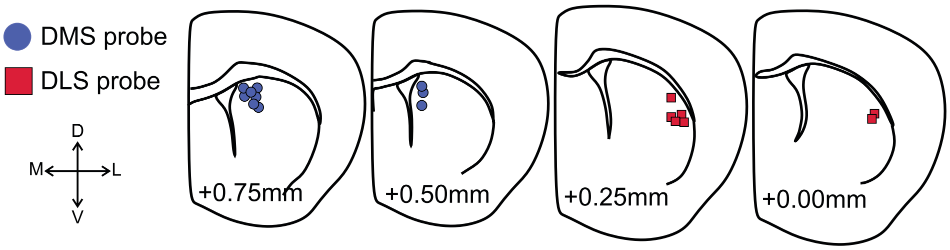


**b** Viral gCaMP8m expression in thalamus

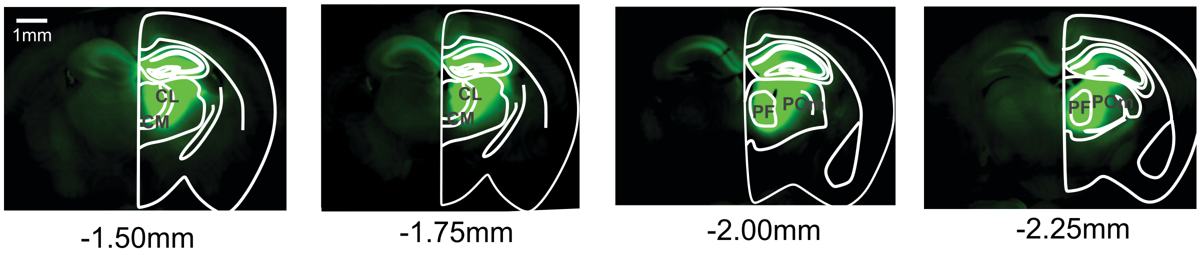


**c** Lack of DMS co-activation in controls

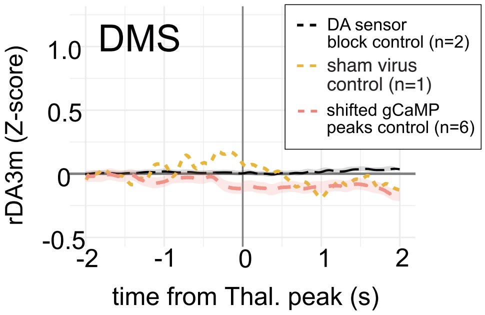

Supplementary Figure 2

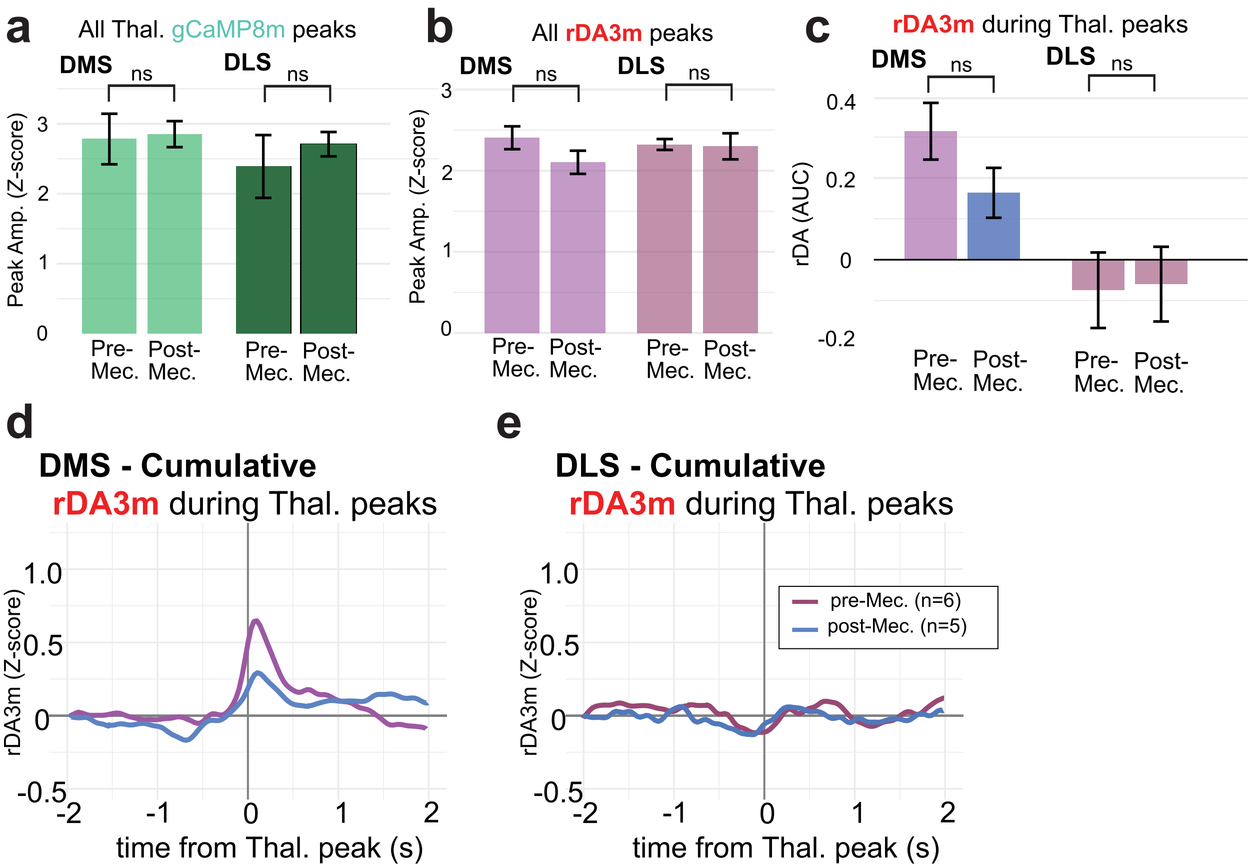

Supplementary Figure 3

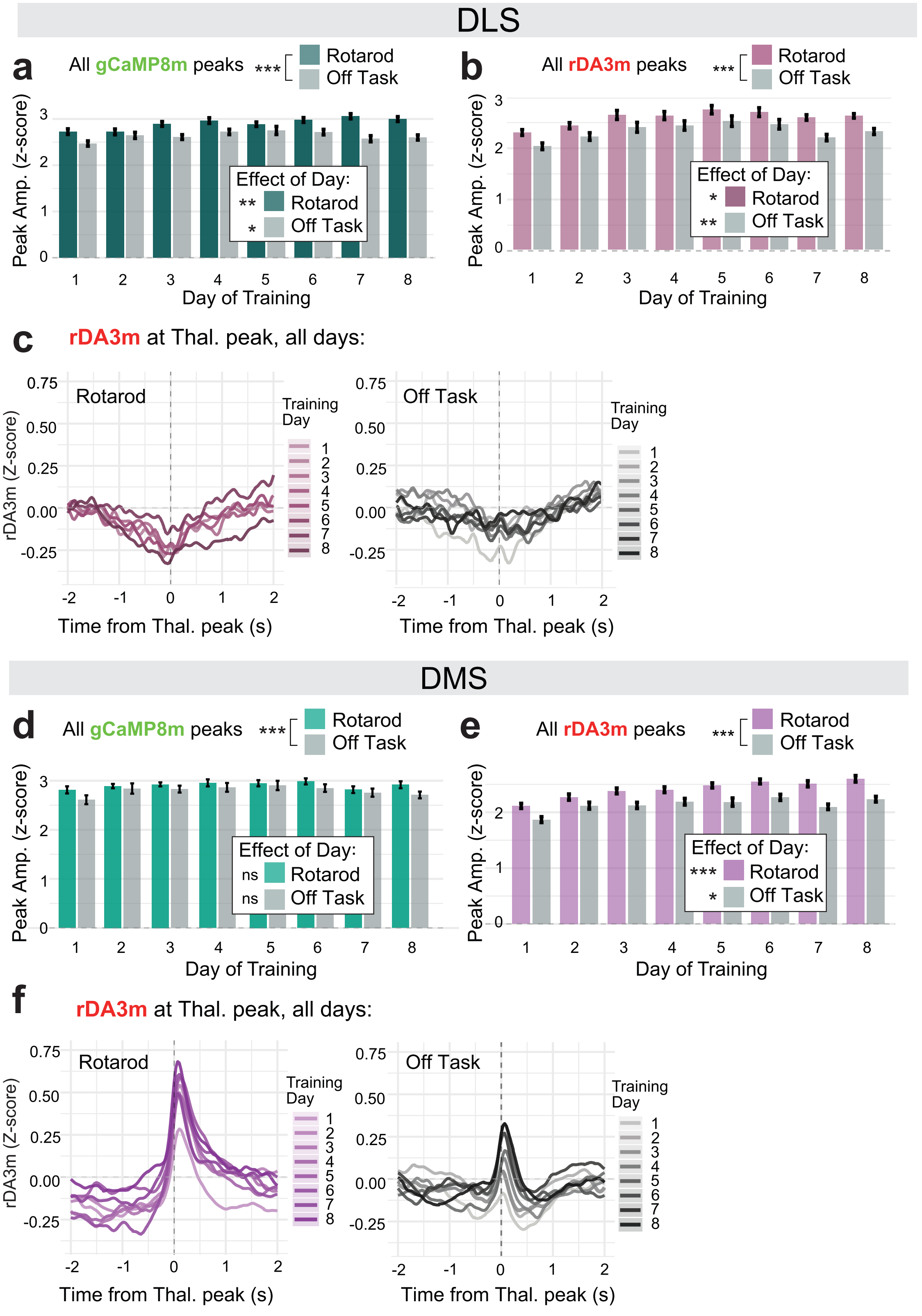

Supplementary Figure 4

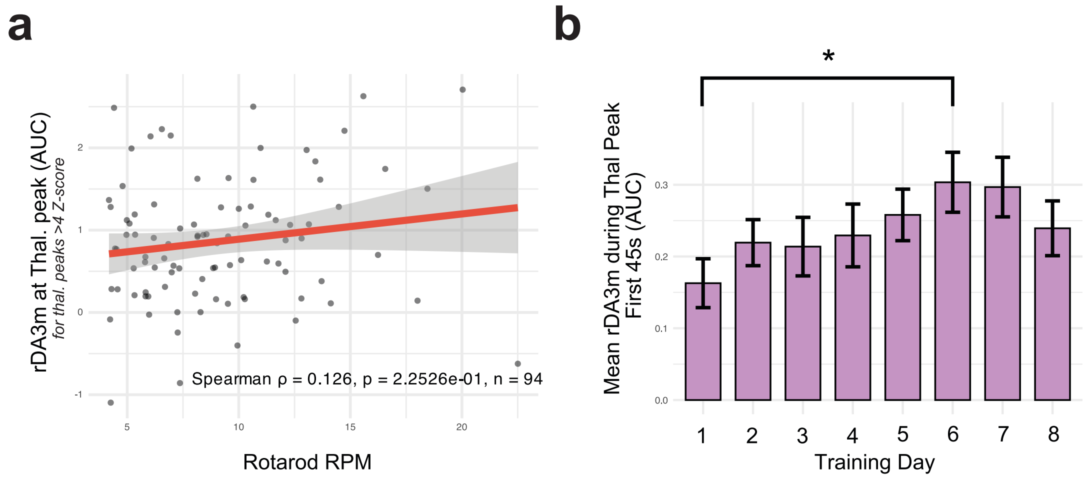

Supplementary Figure 5

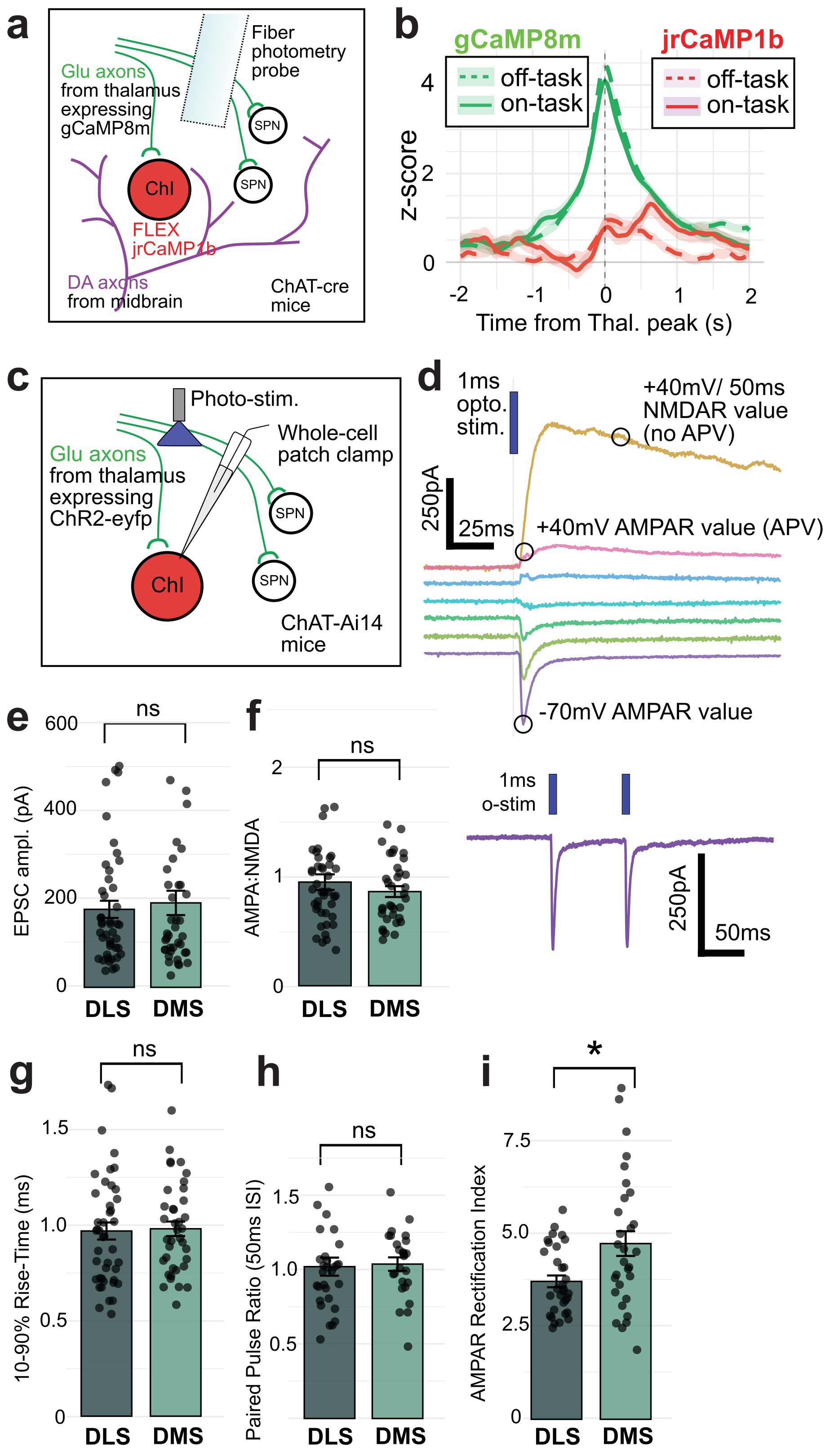

Supplementary Figure 6

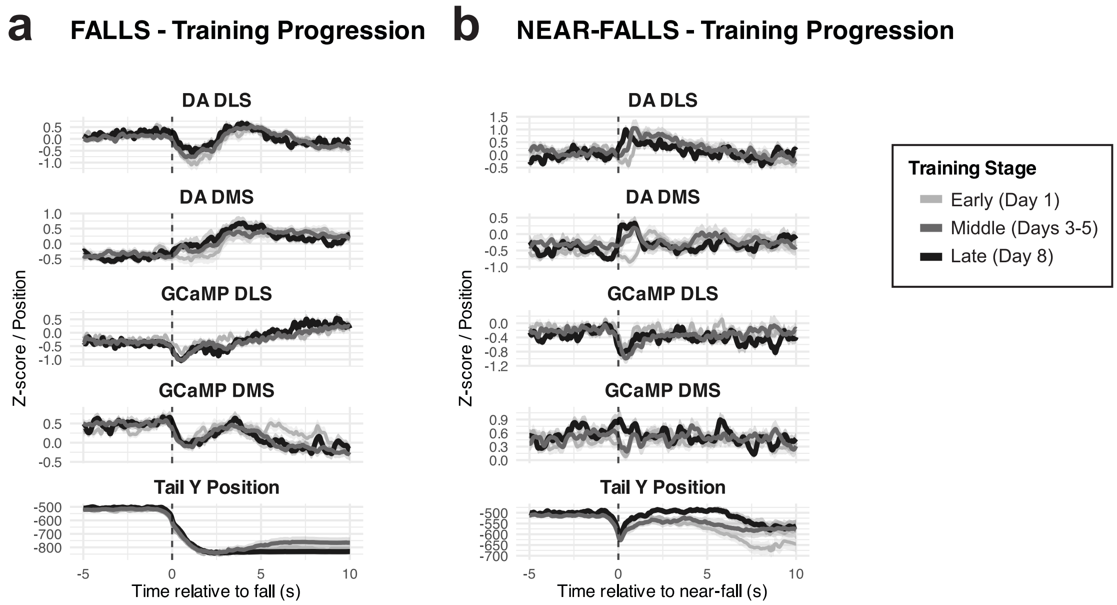

Supplementary Figure 7

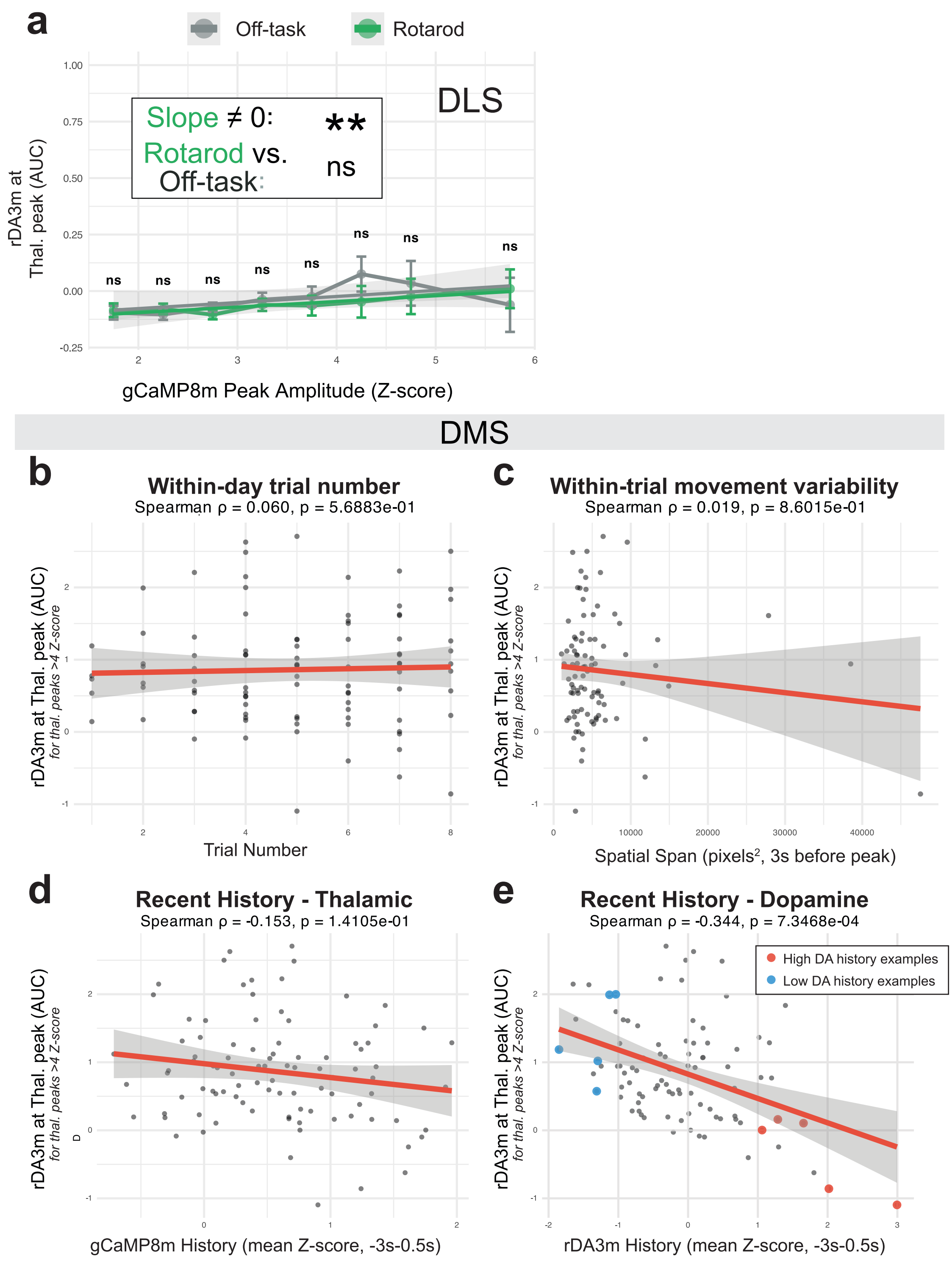
